# Herbivore insect small RNA effector suppress plant defense by cross-kingdom gene silencing

**DOI:** 10.1101/2023.11.18.567654

**Authors:** Wen-Hao Han, Shun-Xia Ji, Feng-Bin Zhang, Hong-Da Song, Jun-Xia Wang, Rui Xie, Xiao-Wei Wang

## Abstract

Herbivore insects deploy salivary effectors to manipulate the defense of their host plants, however, whether insect small RNAs (sRNAs) act as effectors to regulate plant-insect interaction is currently unclear. Here, we report that a microRNA (miR29-b) from the saliva of phloem-feeding insects can transfer into the host plant phloem and fine-tune the host defense. The salivary gland’s abundant miR29-b was produced by insect Dicer 1 and insect salivary exosome is involved in its transferring and releasing into the host plant. Insect miR29-b effector hijacks plant Argonaute 1 to silence host defense gene *Bcl-2-associated athanogene 4* (*BAG4*). Silencing of *BAG4* suppressed the expression of phenylalanine ammonia-lyase and the accumulation of salicylic acid (SA), therefore negatively regulating host defense against herbivore insects. miR29-b is highly conserved in Hemiptera, Coleoptera, Hymenoptera, Orthoptera, and Blattaria insects and also targets the *BAG4* gene. Notably, *BAG4* orthologs exist in a wide range of plant species and may as the target of insect miR29-b. Our work provides new insight into the intriguing defense and counter-defense between herbivores and plants.

**Teaser:** Phloem-feeding insects produce and transfer small RNA into the host plants to fine-tune plant basal defense by cross-kingdom gene silencing.

## Introduction

Herbivore insects have evolved to actively manipulate the defense of their host plants during the 400 million years of co-evolution. Among their armory, saliva effectors stand out as the most prevalent, potent, and ever-evolving weapons utilized in the ongoing arms race with their hosts. Insect saliva effectors have been shown to alter the defense of host plants through various mechanisms (*1–4*). As such, saliva effectors are a new focus in insect-plant interactions and agricultural pest management (*4*, *5*). To date, the known insect saliva effectors have predominantly been proteins and several small molecule compounds capable of disrupting plant immune signaling responses or hindering the production of resistant substances (*1–3*).

Small RNAs (sRNAs) regulate various key biological progress by manipulating gene expression in a highly specific manner (*6*). Typically consisting of 20 to 30 nucleotides, sRNA exerts their influence by complementarily pairing with messenger RNA (mRNA), thereby regulating protein production through post-transcriptional and translational gene silencing (*6*, *7*). MicroRNAs (miRNAs) represent one of the most extensively studied sRNA classes. Remarkably, miRNAs can serve as mobile silencing signals, capable of shuttling between interacting organisms and influencing each other’s gene expression (*8–12*). This phenomenon is known as cross-kingdom RNA interference (*8*, *13*). However, the role of insect miRNAs as cross-kingdom effectors to manipulate plant defense has not been demonstrated, and their target genes in the plant remain unknown.

The whitefly, *Bemisia tabaci* (Gennadius) is a species complex with at least 40 cryptic species, among which *B.tabaci* Middle East-Asia Minor 1 (MEAM1) and Mediterranean (MED) types have invaded more than 100 countries and caused great damage to agriculture production (*14–16*). Whiteflies are extremely polyphagous, with more than 600 species of host plants documented and showing remarkable host adaptability (*17*). As a phloem-feeding herbivore, whiteflies pierce the host epidermis, deftly siphoning nutrient-rich phloem sap through needle-like mandibles, and in this progress, they rely heavily on saliva and saliva effectors to subdue host resistance and facilitate feeding (*1*, *3*, *15*, *18*). A comprehensive and in-depth understanding of how whitefly effectors overcome plant defense mechanisms could help explain its pervasive host adaptability, as well as formulate a sustainable control strategy (*19*, *20*). All the whitefly effectors studied so far are proteins (*15*). Prior research has evidenced the presence of whitefly sRNAs in the tomato phloem (*21*), however, the function of these sRNAs in host plants remains to be established.

Here, we elucidate that miR29-b in whitefly salivary glands acts as an effector, suppressing tobacco defense by hijacking host Argonaute 1 (AGO1). We also uncover the mechanisms underlying *Bt*miR29-b production and release in whiteflies and identify its target, *Bcl-2-associated athanogene 4* (*BAG4*) in host plants. Additionally, we highlight the conservation of miR29-b in phloem-feeding insects and the presence of corresponding *BAG4* genes in various plant species. Notably, this study represents the first report of an insect sRNA effector fine-tuning plant defense through cross-kingdom gene silencing.

## Results

### Whitefly salivary miR29-b is secreted into plants during feeding

Previously, we have identified and analyzed miRNAs in tobacco plants infested by whiteflies (*22*). To identify potential whitefly miRNA effectors, we co-analyzed the miRNAs in tobacco plants and the whitefly miRNA database. *Bt*miR29-b, a known whitefly miRNA, was also detected in whitefly-infested tobacco and was chosen as a possible salivary miRNA effector. The sequenced miRNA from *Bt*miR29-b was 21 nucleotides (nt) in length, consistent with bioinformatic analysis **(fig. S1A)**. The reverse transcription real-time quantitative PCR (RT-qPCR) revealed higher expression of *Bt*miR29-b in the whitefly salivary glands **(Fig. 1A)** and higher levels in adult whiteflies compared to nymphs **(Fig. 1B)**.

**Fig. 1.**
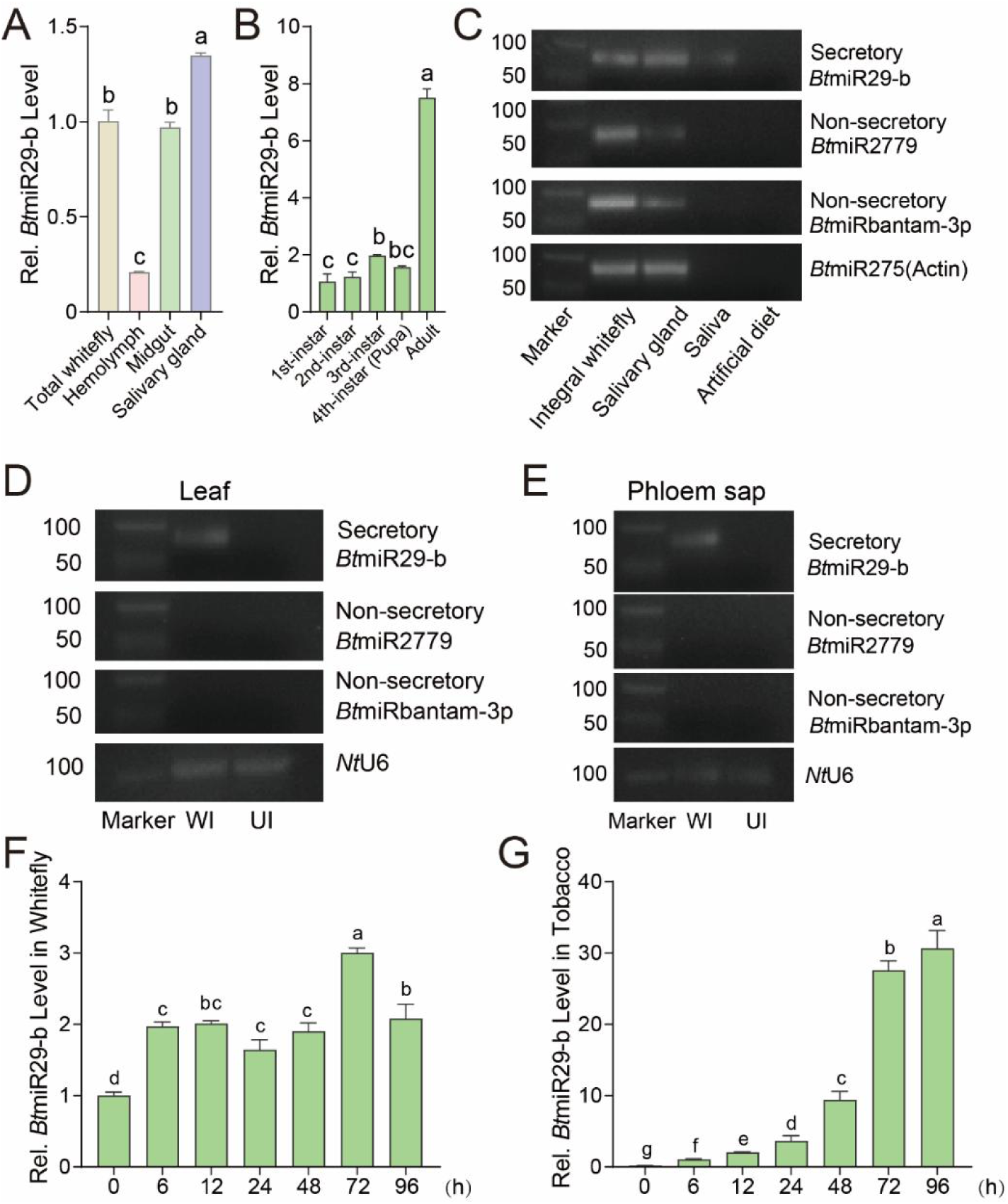
Identification of miRNA effectors released into the host tobacco by whitefly. (**A**) *Bt*miR29-b exhibited higher abundance in the whitefly salivary gland than in other organs. (**B**) *Bt*miR29-b expression varied across different developmental stages of whiteflies. (**C**) Secretory *Bt*miR29-b was detected in whitefly salivary glands and saliva, contrasting with the non-secretory *Bt*miR2779 and *Bt*miRbantam-3p, which were absent in the saliva. (**D, E**) Secretory *Bt*miR29-b was only identified in the whitefly-infested host plant leaf and phloem sap. WI: whitefly infested; UI: uninfested. (**F**) *Bt*miR29-b levels in newly emerged whiteflies differed based on their feeding times on host tobacco (h: Hour). (**G**) *Bt*miR29-b expression in whitefly-infested host tobacco changed across various whitefly feeding times (h: Hour). Values are mean ± SEM; *n* = 3 for **A** (100 whiteflies/1000 organs for each repeat), **B** (100 whiteflies for each repeat), and **F** (300 whiteflies for each repeat); *n* = 6 for **G**. One-way ANOVA followed by Fisher’s least significant difference (LSD) test was used for significant difference analysis. Lowercase letters indicate significant differences between treatments at *P* < 0.05.

We then examined whether *Bt*miR29-b could be secreted into plants during whitefly infestation. Two whitefly miRNAs *Bt*miR2779 and *Bt*miRbantam-3p, which cannot be released into the plant (*21*), were chosen as negative controls. We detected *Bt*miR29-b, *Bt*miR2779, and *Bt*miRbantam-3p in both whitefly tissues and infested tobacco plants by reverse transcription polymerase chain reaction (RT-PCR). The results showed the presence of all three miRNAs in the entire whitefly sample, whereas only *Bt*miR29-b was enriched in the salivary gland and detected in the saliva **(Fig. 1C)**. In addition, *Bt*miR29-b was the only miRNA detected in leaves **(Fig. 1D)** and the phloem exudate of whitefly-infested tobacco plants **(Fig. 1E)**. When newly emerged whiteflies fed on tobacco, the levels of *Bt*miR29-b in the whiteflies upregulated **(Fig. 1F)**, and the abundance of *Bt*miR29-b in host tobacco plants gradually increased **(Fig. 1G)**. We also examined whether *Bt*miR29-b could transfer into other host plants through whitefly feeding. Similarly, *Bt*miR29-b was detected in whitefly-infested Arabidopsis and cotton plants, as well as in their phloem sap **(fig. S1, B and C)**. These results show that *Bt*miR29-b is a secretory miRNA that can be released into different host plants, potentially acting as an effector during whitefly infestation.

### *Bt*miR29-b suppresses host defense against whiteflies

Next, we evaluated whether *Bt*miR29-b affects the host defense against whiteflies. We overexpressed *Bt*miR29-b in tobacco plants using an artificial miRNA vector **(fig. S2, A and B)** and found that whiteflies performed better on *Bt*miR29-b-expressing tobacco plants **(Fig. 2A)**. To further confirm the essential role of the *Bt*miR29-b effector in enhancing whitefly performance on the host, we overexpressed an artificial target mimic of *Bt*miR29-b (amiR29-b) in the host to inhibit its function (*23*, *24*) **(fig. S2C)**. We found that *Bt*miR29-b level in host tobacco overexpressing amiR29-b was reduced **(Fig. 2B)**, and the whiteflies exhibited a reduced survival rate and fecundity on amiR29-b-overexpressing tobacco **(Fig. 2C)**. These results indicate that *Bt*miR29-b may suppress the defense of the tobacco host.

**Fig. 2.**
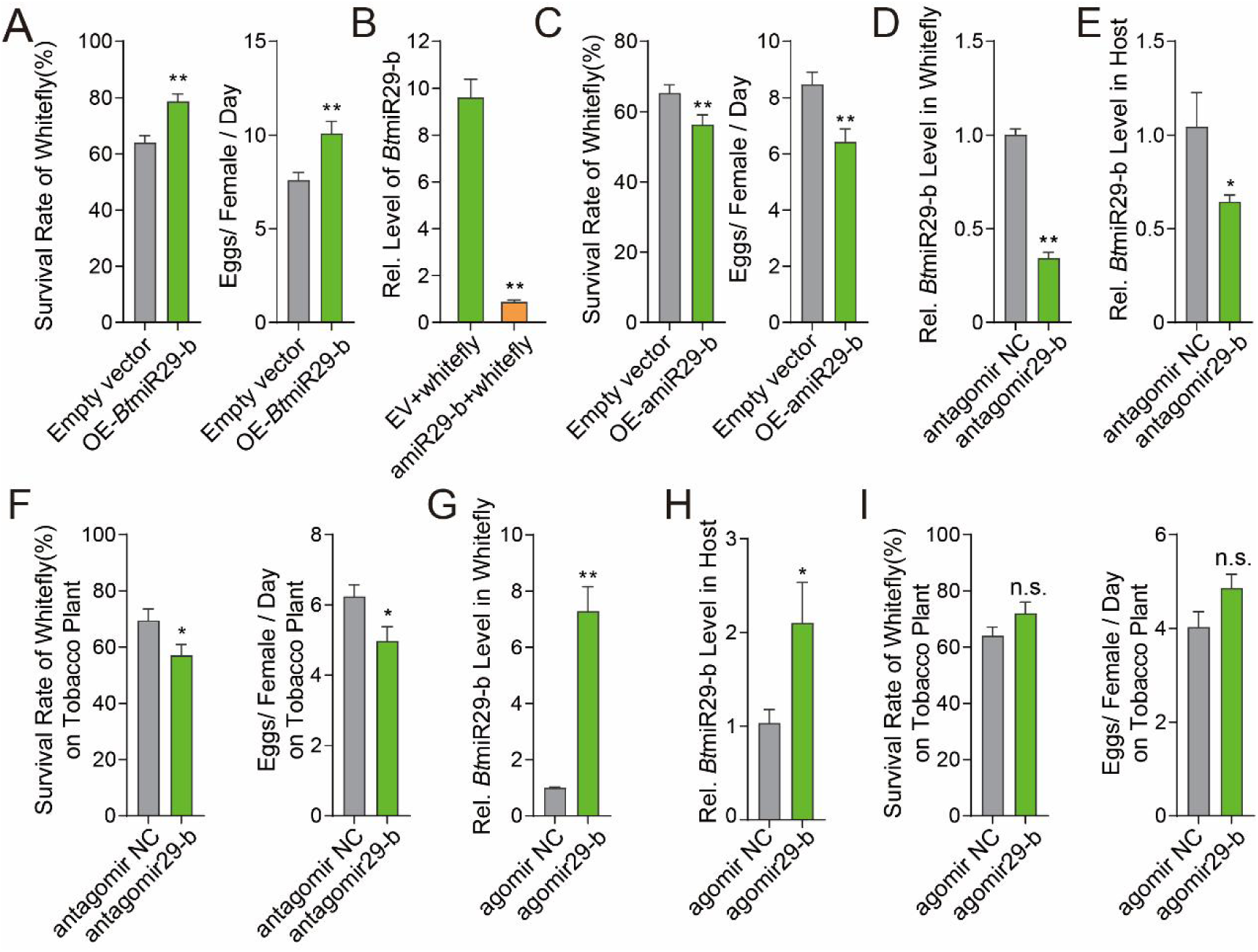
*Bt*miR29-b effector enhances whitefly performance. (**A**) Whiteflies performed better on *Bt*miR29-b-overexpressed tobacco. (**B**) Overexpressing amiR29-b in host tobacco suppressed *Bt*miR29-b accumulation upon whitefly feeding. (**C**) Overexpressing amiR29-b in host tobacco enhanced defense against whiteflies. (**D**) Whitefly *Bt*miR29-b levels were reduced upon antagomir29-b feeding. (**E**) *Bt*miR29-b release into host tobacco decreased when whiteflies fed on antagomir29-b. (**F**) Antagomir29-b-fed whiteflies exhibited poorer performance on tobacco hosts. (**G**) *Bt*miR29-b levels increased significantly in whiteflies upon agomir29-b feeding. (**H**) BtmiR29b released into host tobacco increased when whiteflies fed on agomir29-b. (**I**) Survival rate and fecundity of agomir29-b-fed whiteflies showed an upward trend. Values are mean ± SEM; *n* = 6 for **B**, **D**, **E**, **G**, and **H** (100 whiteflies or 3 tobacco plants for each repeat); *n* = 20 for **A**, **C**, **F,** and **I**. Student’s *t*-test (two-tailed) was used for significant difference analysis. n.s., not significant; *, *P* < 0.05; **, *P* < 0.01.

To investigate whether *Bt*miR29-b is a crucial effector of whiteflies, we employed antagomir, an inhibitor that reduces the number of active miRNAs by pairing with homologous miRNAs, to silence it in whiteflies. The treated whiteflies exhibited a reduced abundance of *Bt*miR29-b **(Fig. 2D)**. Meanwhile, tobacco plants infested by the treated whiteflies showed a lower level of *Bt*miR29-b **(Fig. 2E)**. Furthermore, the survival rate and fecundity of the treated whiteflies significantly decreased when they fed on the host plant **(Fig. 2F)**, but were not affected when fed on an artificial diet **(fig. S2D)**. Next, we overexpressed *Bt*miR29-b in whiteflies by feeding them agomir, a stable miRNA mimic that enhances the expression of miRNAs. Whiteflies fed with agomir29-b had significantly higher levels of intrinsic *Bt*miR29-b **(Fig. 2G)** and released more of it into the host plant **(Fig. 2H)**. Bioassay results showed that, compared with whiteflies fed with the negative control (agomir NC), whiteflies fed with agomir29-b on the tobacco plant displayed an upward trend in survival rate and fecundity when feeding on the host plants **(Fig. 2I)**. No such trend was observed among whiteflies fed an artificial diet **(fig. S2E)**. These results demonstrate that *Bt*miR29-b can suppress the host plant defense to enhance whitefly fitness.

### Whitefly Dicer1 is involved in *Bt*miR29-b production

Dicer proteins contribute to the production of intrinsic miRNAs in insects (*25*, *26*). We supposed that they are also involved in generating miRNA effectors. Sequence analysis revealed that the whitefly genome contains two *Dicer-like* genes, namely *BtDicer1* and *BtDicer2*. To investigate the role of *BtDicer* in *Bt*miR29-b production, we simultaneously silenced *BtDicer1* and *BtDicer2* in whiteflies by feeding them a mixture of ds*BtDicer1*/*2*. The expression of *BtDicer1* and *BtDicer2* was significantly reduced **(Fig. 3A)**, and the abundance of *Bt*miR29-b in the whiteflies **(Fig. 3B)** and the infested host was synchronously reduced **(Fig. 3C)**. Similarly, the performance of ds*BtDicer1*/*2* whiteflies was decreased on the host plant **(fig. S3A)**, but remained unaffected on an artificial diet **(fig. S3B)**.

**Fig. 3.**
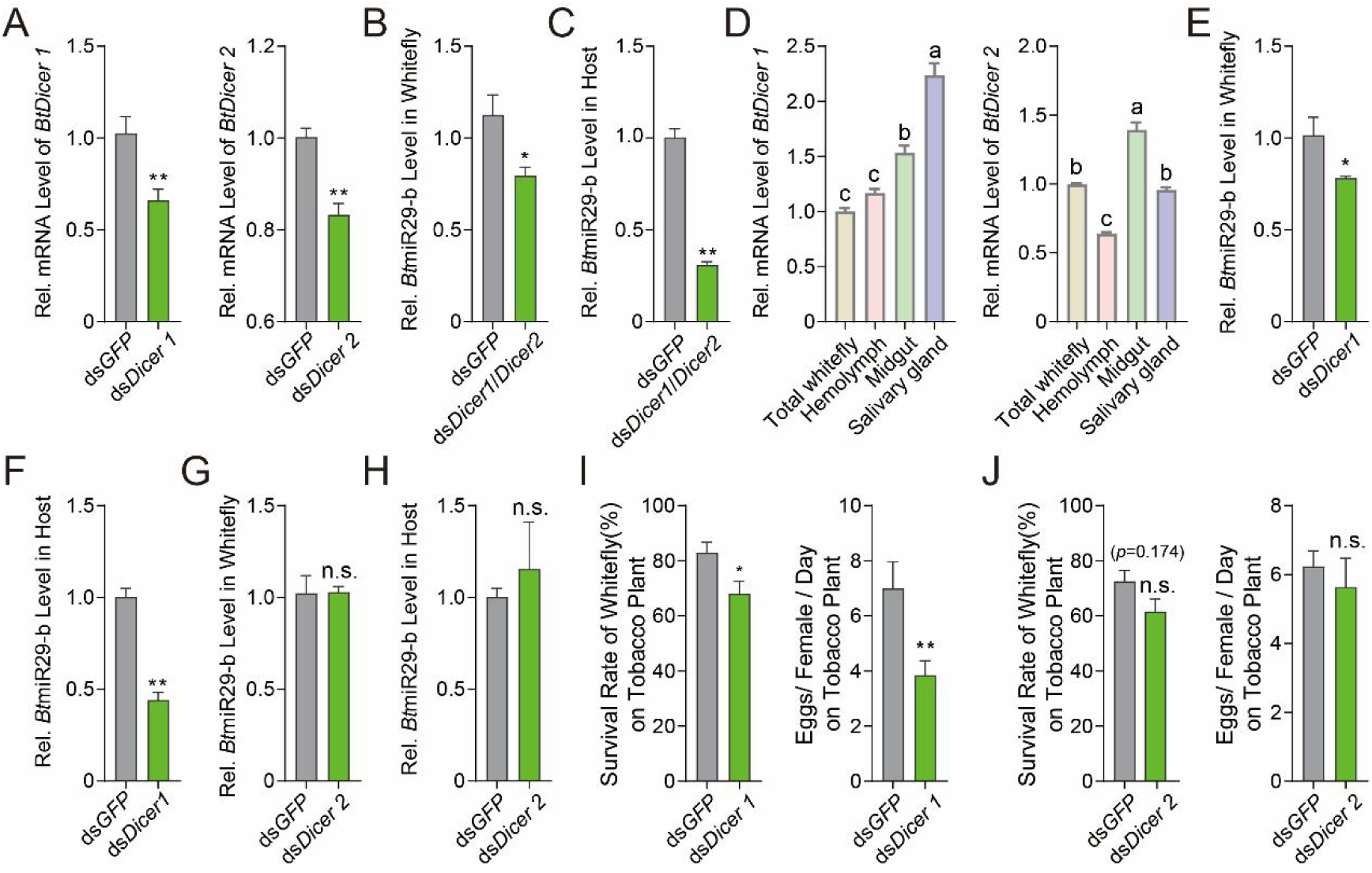
*Bt*Dicer1 mediates *Bt*miR29-b effector production. (**A**) Successful RNA interference (RNAi) of *BtDicer1* and *BtDicer2* through dsRNA feeding. (**B, C**) *Bt*miR29b levels in *BtDicer1*/*BtDicer2*-silenced whiteflies decreased, leading to reduced *Bt*miR29b release into host tobacco during their feeding. (**D**) Expression of *BtDicer1* and *BtDicer2* in total whitefly and various organs. (**E, F**) *Bt*miR29b levels decreased in *BtDicer1*-silenced whiteflies, resulting in less *Bt*miR29b release into host tobacco. (**G, H**) Silencing *BtDicer2* did not affect *Bt*miR29b content or its transfer to host tobacco. (**I**) *BtDicer1*-silenced whiteflies displayed reduced performance on host tobacco. (**J**) *BtDicer2* silencing did not affect whitefly performance on host tobacco. Values are mean ± SEM; *n* = 6 for A, **B**, **C**, **E**, **F**, **G**, and **H** (100 whiteflies or 3 tobacco plants for each repeat); *n* = 3 for **D** (100 whiteflies/ 1000 organs for each repeat); *n* = 20 for **I** and **J**. One-way ANOVA followed by Fisher’s least significant difference (LSD) test or Student’s *t*-test (two-tailed) was used for significant difference analysis. Lowercase letters indicate significant differences between treatments at *P* < 0.05. n.s., not significant; *, *P* < 0.05; **, *P* < 0.01.

Phylogenetic analysis of Dicer proteins in whiteflies and sequence comparison revealed differentiation between *Bt*Dicer1 and *Bt*Dicer2 **(fig. S3C)**. Apparently, only the *BtDicer1* gene exhibited high expression in the salivary glands of whiteflies **(Fig. 3D)**, similar to *Bt*miR29-b **(Fig. 1A)**. We then individually knocked down *BtDicer1* and *BtDicer2*. The abundance of *Bt*miR29-b was suppressed in whiteflies with *BtDicer1* knockdown **(Fig. 3E)**, and tobacco plants infested by whiteflies with *BtDicer1* knockdown showed reduced *Bt*miR29-b content **(Fig. 3F)**. In contrast, these changes were not observed in whiteflies with *BtDicer2* knockdown and in the host plants they infested **(Fig. 3, G and H)**. Interestingly, on tobacco plants, only the performance of whiteflies fed with ds*BtDicer1* significantly decreased **(Fig. 3I)** but not ds*BtDicer2* **(Fig. 3J)**. While the performance of both ds*BtDicer1* and ds*BtDicer2* whiteflies remained unaltered on an artificial diet **(fig. S3, D and E)**. These data indicate that the production of *Bt*miR29-b depends on whitefly Dicer1.

### Whitefly exosome is involved in *Bt*miR29-b secretion into plants

Next, we investigated whether the exosome system of the whitefly was responsible for releasing sRNA effectors from cells into saliva. Triton X-100, a detergent that can rupture exosomes (*10*, *27*, *28*), was added to an artificial diet of whiteflies. Host tobacco infested by whiteflies fed with Triton X-100 displayed reduced *Bt*miR29-b content **(Fig. 4A)**. Subsequently, we explored whether the key genes involved in the whitefly exosome system are responsible for sRNA effector transport. We identified exosome formation genes, *BtCD63* (*29*), *BtSyntenin* (*30*), and *BtSMPD*(*31*), in whiteflies, and observed high expression of all these genes in the midgut **(Fig. 4B)**. When we silenced them by feeding dsRNA, less *Bt*miR29-b was detected in the tobacco **(Fig. 4, C, D and E)**. The performance of ds*BtCD63*, ds*BtSyntenin*, and ds*BtSMPD* treated whiteflies was impacted on tobacco plants **(fig. S4A)**, but not on an artificial diet **(fig. S4B)**. These data indicate that the whitefly exosome system is responsible for releasing sRNA effectors into saliva.

**Fig. 4.**
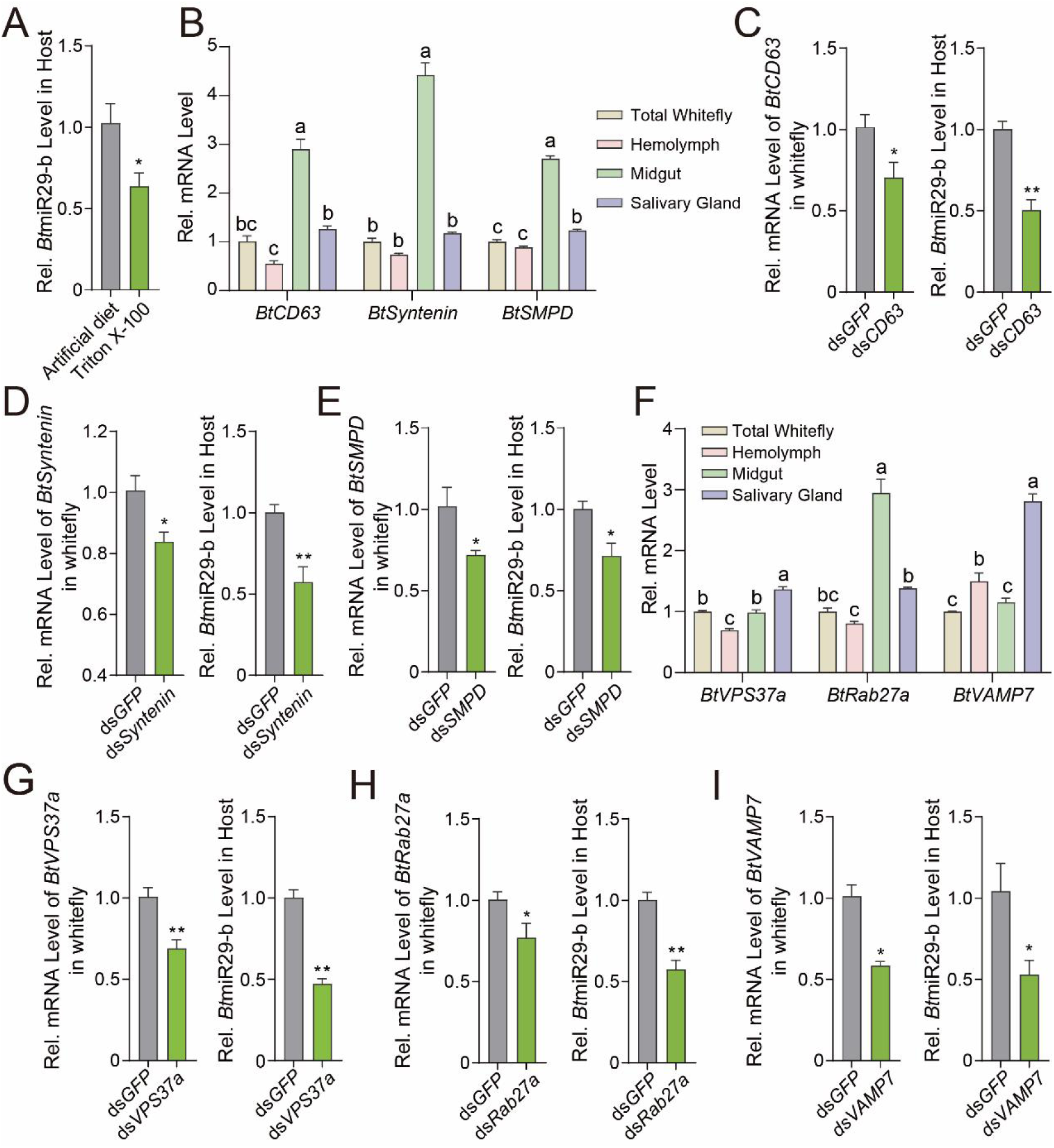
Whitefly exosome systems in *Bt*miRNA29-b secretion. (**A**) *Bt*miR29b release into host tobacco decreased when whiteflies were fed 0.01 mM TritonX-100. (**B**) Expression of *BtCD63*, *BtSyntenin*, and *BtSMPD* genes in whiteflies and different organs. (**C**), (**D**), (**E**) Silencing *BtCD63*, *BtSyntenin*, and *BtSMPD* genes in whiteflies led to reduced *Bt*miR29b release into host tobacco upon infestation. (**F**) Expression of *BtVPS37a*, *BtRab27a*, and *BtVAMP7* genes in whiteflies and different organs. (**G**), (**H**), (**I**) Silencing *BtVPS37a*, *BtRab27a*, and *BtVAMP7* genes in whiteflies led to reduced *Bt*miR29b release into host tobacco upon infestation. Values are mean ± SEM; *n* = 6 for **A**, **C**, **D**, **E**, **G**, **H**, and **I** (100 whiteflies or 3 tobacco plants for each repeat); *n* = 3 for **B** and **F** (100 whiteflies/ 1000 organs for each repeat). One-way ANOVA followed by Fisher’s least significant difference (LSD) test or Student’s *t*-test (two-tailed) was used for significant difference analysis. Lowercase letters indicate significant differences between treatments at *P* < 0.05. n.s., not significant; *, *P* < 0.05; **, *P* < 0.01.

Furthermore, we examined the role of whitefly VPS37a, involved in cargo sorting (*32*); Rab27a, involved in multivesicular bodies (MVB) transport (*33*); and VAMP7 (*34*), involved in *Bt*miR29-b secretion. *BtRab27a* exhibited high expression in both the midgut and salivary glands, while *BtVPS37a* and *BtVAMP7* were highly expressed in the salivary glands **(Fig. 4F)**. We found that, when *BtVPS37a*, *BtRab27a*, and *BtVAMP7* were silenced by feeding dsRNA, less *Bt*miR29-b was released into the host tobacco plant upon whitefly infestation **(Fig. 4, G, H, and I)**. Whiteflies with *VPS37a*, *Rab27a*, and *VAMP7* knockdown performed worse on the host plant but were not affected when fed on an artificial diet **(fig. S4, C to H)**. These results demonstrate that VPS37a, Rab27a, and VAMP7 play a role during the secretion of whitefly sRNA effector into the host.

### *Bt*miR29-b targets the plant defense gene *BAG4*

To investigate how *Bt*miR29-b alters tobacco defense against whiteflies, we predicted *Bt*miR29-b targets in tobacco hosts using Target Finder (https://github.com/carringtonlab/TargetFinder). Our analyses revealed that *Bt*miR29-b targets the transcripts of *BAG4* and several other genes in tobacco plants **(Table S1)**. Firstly, we examined the transcript level of *NtBAG4* after whitefly infestation. The results showed that whitefly infestation repressed the *NtBAG4* gene **(Fig. 5A)**. Additionally, plants infested by whiteflies fed with *Bt*miR29-b antagomir29-b showed enhanced *NtBAG4* expression **(Fig. 5B)**, while those fed with agomir29-b repressed its expression **(Fig. 5C)**. Meanwhile, *NtBAG4* was also inhibited in tobacco plants that artificially expressed *Bt*miR29-b **(Fig. 5D)**.

**Fig. 5.**
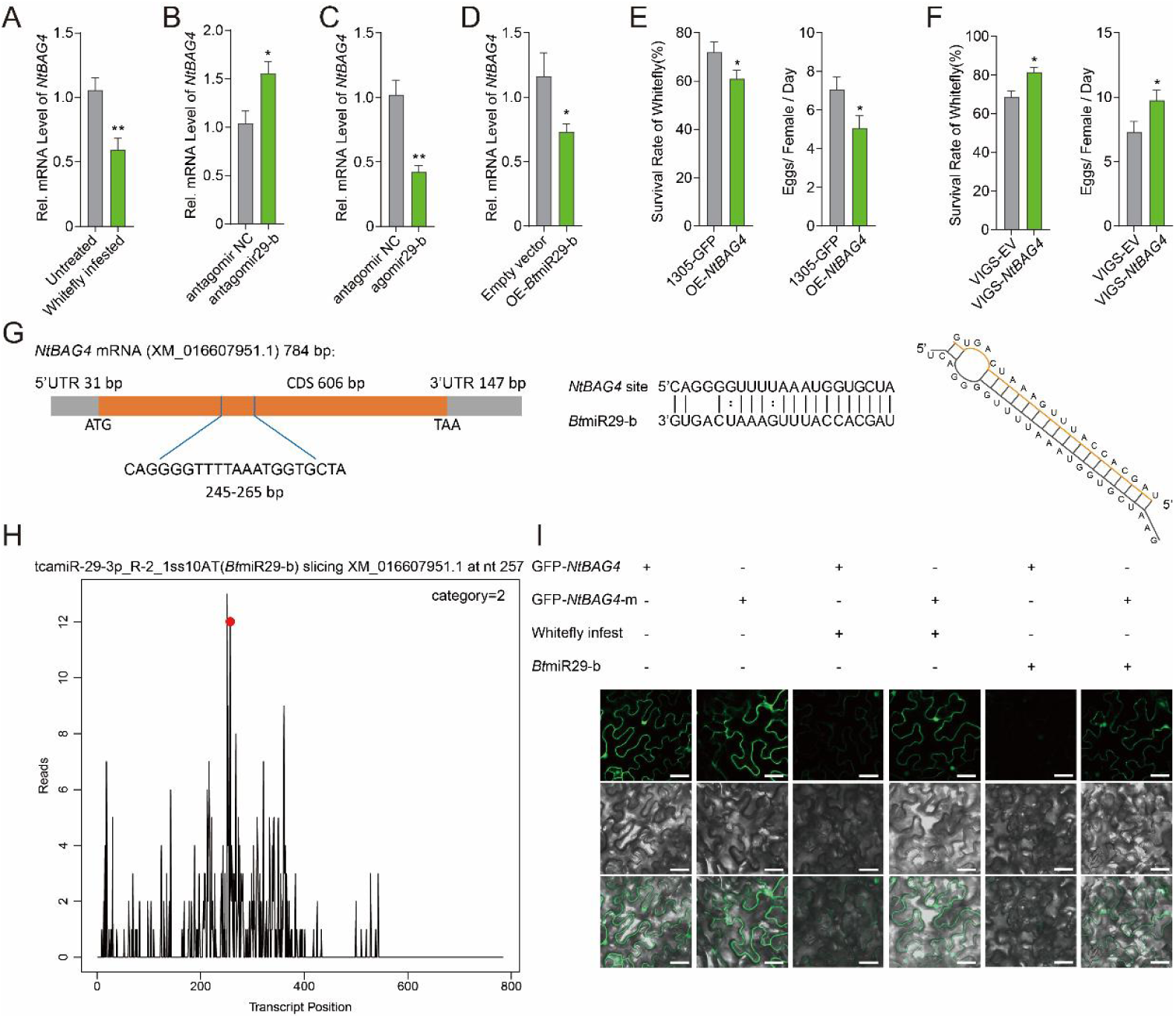
*Bt*miR29-b suppresses tobacco gene *NtBAG4* and inhibits host defense. (**A**) Whitefly infestation suppressed the tobacco target gene *NtBAG4* by *Bt*miR29-b. (**B**) The *NtBAG4* mRNA level significantly increased in tobacco infested by antagomir29-b-fed whiteflies. (**C**) The *NtBAG4* mRNA level was lower in tobacco infested by agomir29-b-fed whiteflies. (**D**) *Bt*miR29-b in tobacco suppressed target the *NtBAG4*. (**E**) Whiteflies performed poorly on *NtBAG4*-overexpressed tobacco. (**F**) Whiteflies thrived on *NtBAG4*-silenced tobacco. (**G**) Schematic illustrating *Bt*miR29-b’s targeting of tobacco *NtBAG4* mRNA. CDS: Coding sequence; UTR: Untranslated regions; ATG: Initiation codon; TAA: Termination codon. (**H**) Target plots (t-plots) of *Bt*miR29-b and its target *NtBAG4* mRNA, validated via degradome sequencing. The red dot indicates the cleavage site analyzed by GSTAr (1.0) cleavage site prediction software, showing the *NtBAG4* transcript was silenced at 257 bp. (**I**) Confocal images displayed GFP-*NtBAG4* and GFP-*NtBAG4*-m expression in *N. benthamiana* during whitefly infestation and *Bt*miR29-b overexpression. Top, GFP. Middle, Bright-field. Bottom, GFP/bright-field overlay. Scale bars, 40 μm. Values are mean ± SEM; *n* = 6 for **A**, **B**, **C**, and **D** (100 whiteflies or 3 tobacco plants for each repeat), *n* = 20 for **E** and **F**; Student’s *t*-test (two-tailed) was used for significance. *, *P* < 0.05; **, *P* < 0.01.

Subsequently, we investigated whether *NtBAG4* is involved in tobacco defense against whiteflies. Overexpression of *NtBAG4* in tobacco **(fig. S5A)** significantly enhanced the host defense against whiteflies **(Fig. 5E)**. Additionally, suppressing the expression of *NtBAG4* by virus-induced gene silencing (VIGS) **(fig. S5B)** benefited whitefly survival on tobacco hosts **(Fig. 5F)**. These results show that the whitefly miRNA effector *Bt*miR29-b can silence the tobacco defense gene *NtBAG4*, thereby enhancing its performance on tobacco.

### *Bt*miR29-b targets the coding region (245 to 256 bp) of *NtBAG4*

Sequence analysis indicated that *Bt*miR29-b targeted the coding region of *NtBAG4* (245 to 256 bp) **(Fig. 5G)**. To determine the cleavage site of tobacco *NtBAG4* by whitefly *Bt*miR29-b, we conducted degradome sequencing of whitefly-infested tobacco samples. Target plots validated that tobacco *NtBAG4* was indeed cleaved by *Bt*miR29-b upon whitefly feeding. The red dot indicates the cleavage site according to the analysis of degradome sequencing, which corresponds to the predicted *Bt*miR29-b target site **(Fig. 5H)**.

To further confirm this finding, we cloned green fluorescent protein (GFP)-tagged full-length *NtBAG4* (GFP-*NtBAG4*) into an overexpression cassette and expressed it in *Nicotiana benthamiana* leaves **(fig S6A)**. We observed the suppression of GFP-tagged *NtBAG4* two days after whitefly infestation **(Fig. 5I)**. However when the *Bt*miR29-b target site of full-length *NtBAG4* was mutated (GFP-*NtBAG4*-m) **(fig S6B)**, whitefly infestation failed to suppress its expression **(Fig. 5I)**. Meanwhile, the expression of GFP-tagged *NtBAG4* was repressed when coexpressed with *Bt*miR29-b **(Fig. 5I)**, while that of mutant *NtBAG4* was not affected **(Fig. 5I)**. These results demonstrate that only the wild-type *NtBAG4* gene was suppressed after whitefly infestation or coexpression of *Bt*miR29-b. In addition, we cloned the GFP-tagged *Bt*miR29-b target site (5’CAGGGGUUUUAAAUGGUGCUA3’) and mutant site into the overexpression cassette **(fig. S6C)**. We observed the suppression of the GFP-tagged *Bt*miR29-b target site by whitefly infestation and coexpression with *Bt*miR29-b, but not in GFP-tagged mutant target sites (**fig. S6D**), further determined that the site we identified in *BAG4* can be targeted by *Bt*miR29-b.

### *Bt*miR29-b silences *NtBAG4* in the host phloem

GUS tags enable the spatial visualization of gene expression and degradation, thus, we further employed this method to investigate where the cleavage of *BAG4* by whitefly *Bt*miR29-b occurs in plant leaves. First, we conducted the coexpression test in the *Nicotiana tabacum* plant by examining the GUS-tagged target expression (**fig. S7A**). Similarly, the expression of GUS-tagged *NtBAG4* and the GUS-tagged *Bt*miR29-b target site was suppressed during *Bt*miR29-b over-expression, while that of GUS-tagged mutant *NtBAG4* or mutant *Bt*miR29-b target site was not affected (**fig. S7B**). Spatial visualization of the GUS driven by the promoter of *NtBAG4* showed that the *NtBAG4* gene was expressed in the phloem, similar to the promoter of *AtSUC2* (sucrose transporters 2), a phloem-specific expression promoter. **(fig. S7C)**. The GUS-tagged *Bt*miR29-b target site driven by the promoter of *NtBAG4* is also expressed in the phloem, but this expression was repressed upon whitefly infestation. Similarly, when *Bt*miR29-b was expressed in the phloem driven by the *AtSUC*2 promoter, the expression of the GUS-tag was also repressed **(fig. S7D)**. These results suggest that *Bt*miR29-b might target the host phloem-expressed defense gene *NtBAG4*.

### *Bt*mi29-b hijacks host AGO1 to suppress host *BAG4* gene

We further investigated how insect sRNA silences the host *BAG4* gene. In plants, miRNAs can be loaded into AGO1 to form the RNA-induced silencing complex (*35*) and then cleavage the transcript of target genes (*36*). We sought to investigate whether whitefly effectors can hijack the plant’s gene-silencing mechanism to target *BAG4*. Firstly, we hypothesized that if host AGO1 is essential for whitefly sRNA effectors to silence host genes, then interfering with the expression of host *AGO1* will enhance plant resistance. In tobacco plants, four *NtAGO1* genes (LOC107806342; LOC107790785; LOC107771391; LOC107789474) have been annotated, and we simultaneously repressed them using VIGS. We found that the VIGS-*NtAGO1*s tobacco plants showed enhanced defense against whiteflies **(fig. S8A)**. Then, we found that, in the VIGS-*NtAGO1*s plant, the *NtBAG4* transcripts and the expression of GFP fused with the *NtBAG4* target site were not suppressed during whitefly infestation and *Bt*miR29-b overexpression **(fig. S8, B and C)**.

To further confirm this result, we obtained the *ago1* mutant Arabidopsis. The *ago1* mutant plants showed enhanced defense against whiteflies **(fig. S8D)**, suggesting that these *Bt*sRNAs could no longer suppress host defense genes. In addition, whitefly infestation reduced the expression of the anti-whitefly gene *AtBAG4* in wild-type plants, but not in *ago1* mutant plants **(fig. S8F)**. These results demonstrate that *Bt*miR29-b may directly hijack AGO1 and cleavage the defense gene *BAG4*.

### *NtBAG4* promotes SA accumulation by upregulating *PAL*

Next, we explored how *NtBAG4*, the target gene of *Bt*miR29-b, positively regulates plant resistance against whitefly. SA is known as a defense phytohormone in plants against phloem-feeding insects (*37*). We observed that tobacco plants overexpressing the *NtBAG4* gene accumulate more intrinsic SA compared to the control **(Fig. 6A)**. SA biosynthesis occurs through the isochorismate synthase (ICS) and phenylalanine ammonia-lyase (PAL) pathways **(Fig. 6B)**. Thus, we explored the mechanism of BAG4-mediated SA accumulation by monitoring the expression of SA synthesizing genes. Quantitative PCR results showed that the expression of key synthetic genes in the ICS pathways, including *NtICS1*, *NtEDS5* (enhanced disease susceptibility), and *NtEPS1* (enhanced pseudomonas susceptibility) was repressed, and *NtICS2* was not affected **(Fig. 6C)**. Notably, the expression of four vital *NtPAL* genes in PAL pathways was enhanced by *NtBAG4* overexpression **(Fig. 6C)**, and maker genes downstream of SA signaling, *NtNPR1* (nonexpresser of PR genes 1), and *NtPR1* (pathogenesis-related gene 1) were also induced **(Fig. 6C)**. Consistently, in tobacco plants with VIGS-*NtBAG4*, the expression of *NtPAL2* was repressed **(Fig. 6D)**.

**Fig. 6.**
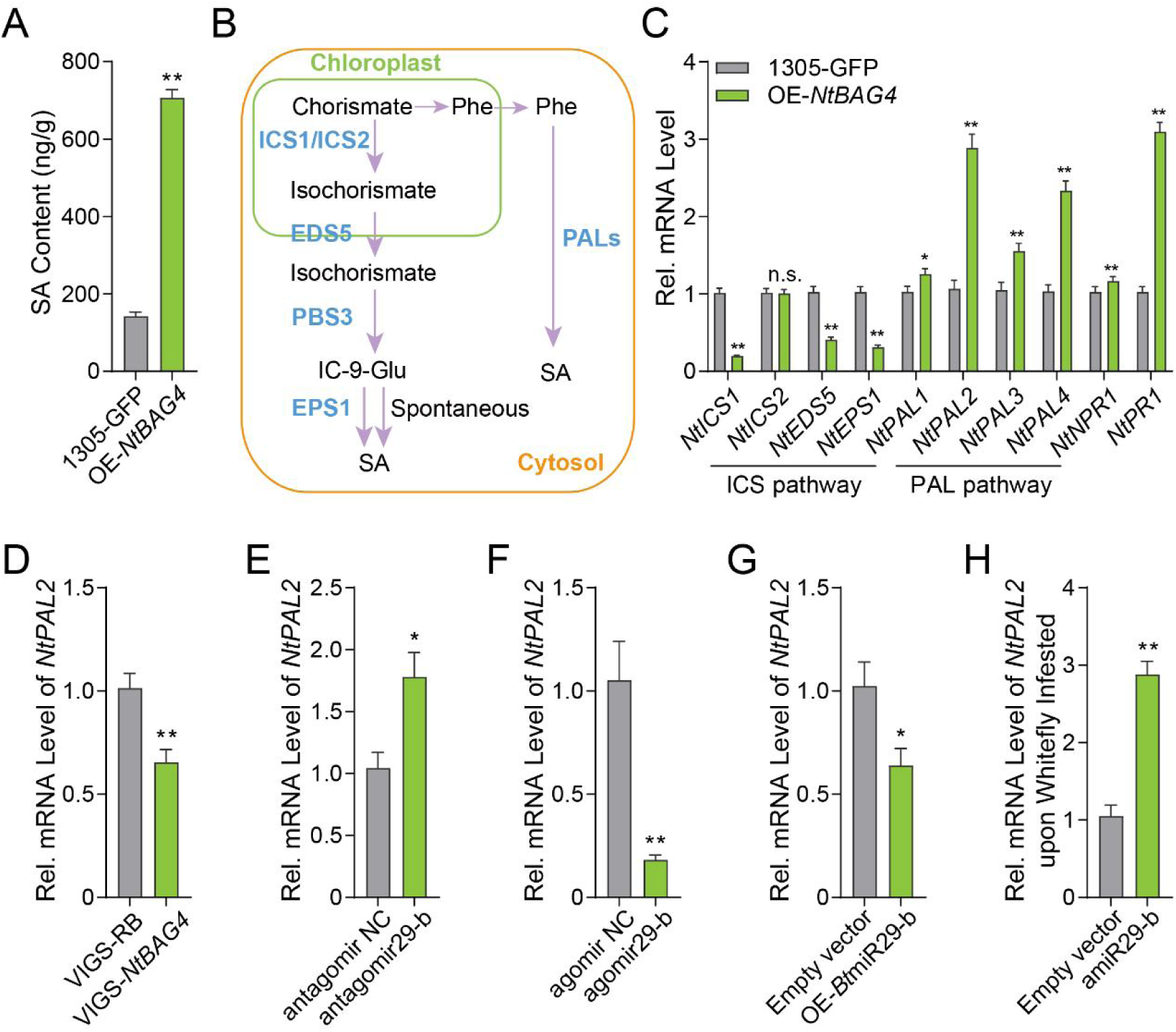
*NtBAG4*’s role in plant SA accumulation. (**A**) *NtBAG4*-overexpressed tobacco displays higher SA accumulation. (**B**) SA biosynthesis pathway in the plant. ICS1: Isochorismate synthase; EDS5: Enhanced Disease Susceptibility 5; PBS3: AvrPphB Susceptible 3; EPS1: Enhanced *Pseudomonas* Susceptibility 1; PAL: Phenylalanine ammonia-lyase. (**C**) *NtBAG4* overexpression suppresses key enzyme genes in the ICS pathway of SA synthesis, induces genes in the PAL pathway, and increases SA marker genes. (**D**) *NtPAL2* mRNA levels decrease in *NtBAG4*-silenced tobacco. (**E**, **F**) *NtPAL2* mRNA was higher in tobacco infested by antagomir29-b-fed whiteflies and lower in tobacco infested by agomir29-b-fed whiteflies. (**G**) *Bt*miR29-b overexpression inhibits *NtPAL2* expression in tobacco. (**H**) amiR29-b overexpression enhances *NtPAL2* mRNA levels in tobacco upon whitefly infestation. Values are mean ± SEM; *n* = 6 (3 tobacco plants for each repeat). Student’s *t*-test (two-tailed) was used for significant difference analysis. n.s., not significant; *, *P* < 0.05; **, *P* < 0.01.

Since *NtBAG4* is associated with *NtPAL* transcript accumulation, we further examined the relationship between *NtPAL* and the whitefly *Bt*miR29-b effector. Whiteflies fed with antagomir29-b were shown to have less *Bt*miR29-b **(Fig. 2D)**, and the level of *NtPAL2* transcripts in tobacco infected with them was significantly increased **(Fig. 6E)**. In contrast, whiteflies fed with agomir29-b had more *Bt*miR29-b **(Fig. 2G)** and significantly repressed *NtPAL2* when infesting tobacco **(Fig. 6F)**. Additionally, tobacco over-expressing *Bt*miR29-b showed reduced expression of *NtPAL2* **(Fig. 6G)**, while the amiR29-b expressed tobacco plant showed higher *NtPAL2* expression upon whitefly infestation **(Fig. 6H)**. These results collectively suggest that BAG4 may promote the accumulation of SA by inducing the expression of key genes in the PAL pathway of SA biosynthesis, and *Bt*miR29-b can interfere with the expression of *NtPAL2* by degrading *NtBAG4*.

### Whitefly Dicer1 and exosome are critical for the suppression of *NtBAG4* and ***NtPAL2* in host plants**

We further determined the essential role of Dicer and exosome in whiteflies for the suppression of defense genes *NtBAG4* and *NtPAL2* in host plants. We evaluated the effect of silencing *Dicer* and exosome-associated genes on the ability of whiteflies to inhibit plant *BAG4* and its downstream *PAL2* gene. The *NtBAG4* mRNA level in tobacco infested by *BtDicer1*-silenced whiteflies was significantly increased compared with control whiteflies, while that of *BtDicer2*-silenced whiteflies was not changed **(fig. S9, A and B)**. Similarly, the expression of the GFP fused to the *NtBAG4* target site was still silenced upon infestation by *BtDicer2*-silenced whiteflies, while the suppression was abolished when fed by *BtDicer1*-silenced whiteflies **(fig. S9C)**. Consistent with this, the mRNA level of *NtPAL2* in tobacco infested by *BtDicer1*-silenced but not *BtDicer2*-silenced whiteflies was significantly increased **(fig. S9. D and E)**.

Next, we found that the *NtBAG4* mRNA level in tobacco fed by Triton X-100 feeding and exosome formation-related genes *BtCD63*/*BtSyntenin*/*BtSMPD*-silenced was significantly increased **(fig. S10, A to D)** and the suppression of the GFP fused to the *NtBAG4* target site was abolished when fed by these treated whiteflies **(fig. S10E)**. Meanwhile, the *NtPAL2* mRNA level was also significantly enhanced in tobacco fed by these whiteflies compared with that fed by control whiteflies **(fig. S10, F to I)**. In addition, when whitefly genes associated with exosome cargo sorting (*BtVPS37a*), transport (*BtRab37a*), and releasing (*BtVAMP7*) were silenced in whiteflies, respectively, treated whiteflies could not suppress *NtBAG4* **(fig. S10, J to M)** and *NtPAL2* **(fig. S10, N to P)**. These results collectively suggest the importance of genes related to miRNA effector production and release in the suppression of host resistance genes.

### *Bt*miR29-b is a conserved sRNA effector in herbivore insects

By analyzing the reported miRNA database (https://www.mirbase.org/), we found that *Bt*miR29-b exists in many Hemiptera, Coleoptera, Hymenoptera, Orthoptera, and Blattaria insect species **(fig. S11A)**. We then cloned it from several Hemiptera pests **(fig. S11B)**, and the sequences showed that *Mp*miR29-b (from the green peach aphid *Myzus persicae*) was consistent with the first 20bp of *Bt*miR29-b **(fig. S11C)**. *Mp*miR29-b was also detected in aphid salivary glands and saliva **(Fig. 7A)**, and in aphid-infested tobacco plants and phloem sap **(fig. S11D and Fig. 7B)**. The aphids showed better performance on tobacco plants overexpressing miR29-b **(Fig. 7C)**, but worse on tobacco plants overexpressing amiR29-b **(Fig. 7D)**. Aphid infestation can also suppress *NtBAG4* **(Fig. 7E)**, which was involved in tobacco defense against aphids **(Fig. 7F)**. These results indicate that miR29-b may function as an sRNA effector and alter plant defense in different Hemiptera insects.

**Fig. 7.**
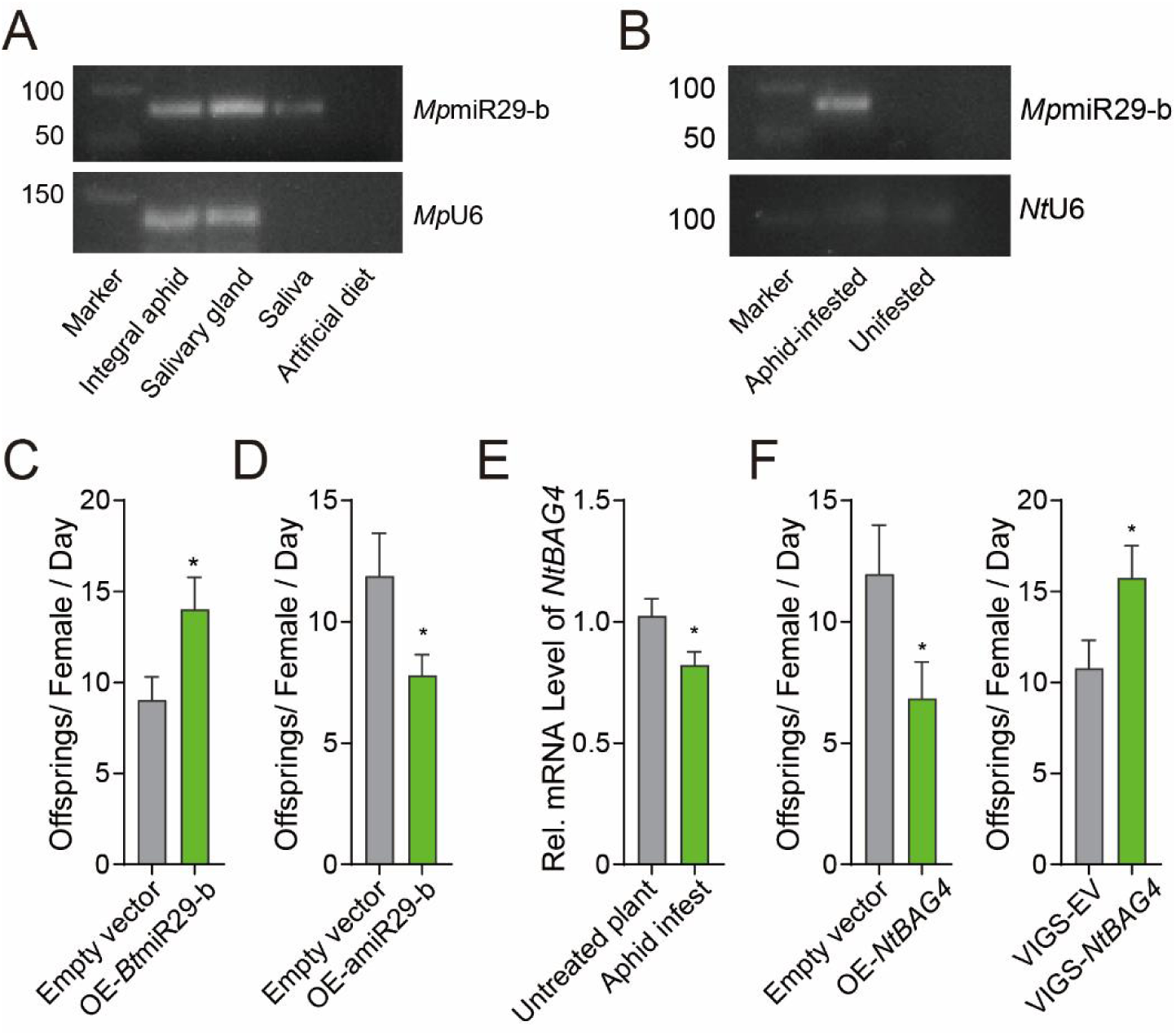
*Bt*miR29-b is a conserved sRNA effector in herbivore insect and play the same role in plant-aphid interaction. (**A**) *Mp*miR29-b was detected in the whitefly salivary gland and saliva. (**B**) *Mp*miR29-b was detected in aphid-infested host phloem sap. (**C**) Aphids perform better on *Bt*miR29-b expressed tobacco. (**D**) Aphids perform worse on amiR29-b expressed tobacco. (**E**) Aphid infestation suppresses *NtBAG4* gene expression. (**F**) *NtBAG4* is involved in tobacco defense against aphids. Values are mean ± SEM; *n* = 20 for **C**, **D**, and **F**; *n* = 12 for **E** (3 tobacco plants for each repeat). Student’s *t*-test (two-tailed) was used for significant difference analysis. *, *P* < 0.05.

### MiR29-b effector targets *BAG4* in a wide range of angiosperm species

We further expanded our analysis of *BAG4* genes to other angiosperm species, with a focus on economically significant crop plants susceptible to phloem-feeding insects. *BAG4* genes are present in all species studied, but their sequences display variations among different plant families **(Table S2)**. We further analyzed potential miR29-b target sites in plants from Solanaceae, Brassicaceae, Malvaceae, Leguminosae, Cucurbitaceae, and Euphorbiaceae families. Although the sequence of *BAG4* transcripts is different in several plant species, the possible target sites can be predicted and shown by Target Finder **(fig. S12)**. We found that the target sites were different among family varieties but conserved within families **(fig. S12)**. Interestingly, computational analyses showed that, in Solanaceae plants, the miR29-b target site within the *BAG4* gene is shared among multiple *Nicotiana* species but differs from that of other species, including tomato **(fig. S12)**. In several pepper and potato plants, the predicted target site of miR29-b in the *BAG4* genes is the same as that of the tomato **(fig. S12)**. These findings collectively suggest that herbivore insect miR29-b effectors may interact with diverse *BAG4* sites in angiosperm species, utilizing distinct binding mechanisms.

### MiR29-b effector targets tomato *BAG4* sites as well

To explore whether miRNA effectors can target different sites to degrade *BAG* in different plant species, we selected tomato plants for further verification. Firstly, we determined that *Bt*miR29-b could also suppress tomato defense against whiteflies **(Fig. 8A)**, indicating that whiteflies suppress tomato defense by miR29-b effector. Then, *SlBAG4* was proved to be positively involved in tomato resistance against whiteflies, as examined by overexpression and TRV-mediated gene silencing **(Fig. 8, B and C; fig. S13, A and B)**, showing that *SlBAG4* may be a potential target for whiteflies to alter tomato defense. Next, we found that whitefly infestation and *Bt*miR29-b expression can also repress *SlBAG4* **(Fig. 8, D and E)**. Meanwhile, we verified the analyzed target site of miR29-b on *SlBAG4* **(fig. S12)** by evaluating the expression of GFP fused with the *SlBAG4* target site, and the results show that whitefly infestation, and *Bt*miR29-b expression cleavage *SlBAG4* site, but not the mutant site **(fig. S13 C and Fig. 8F)**. Furthermore, the overexpression of amiR29-b suppressed *Bt*miR29-b accumulation in tomatoes and increased the mRNA level of *SlBAG4* upon whitefly infestation **(fig. S13, D and E; Fig. 8G)** and enhances tomato defense against whiteflies **(Fig. 8H)**. These results collectively reveal that whitefly *Bt*miR29-b can also degrade the tomato *BAG4* gene and suppress tomato defense responses.

**Fig. 8.**
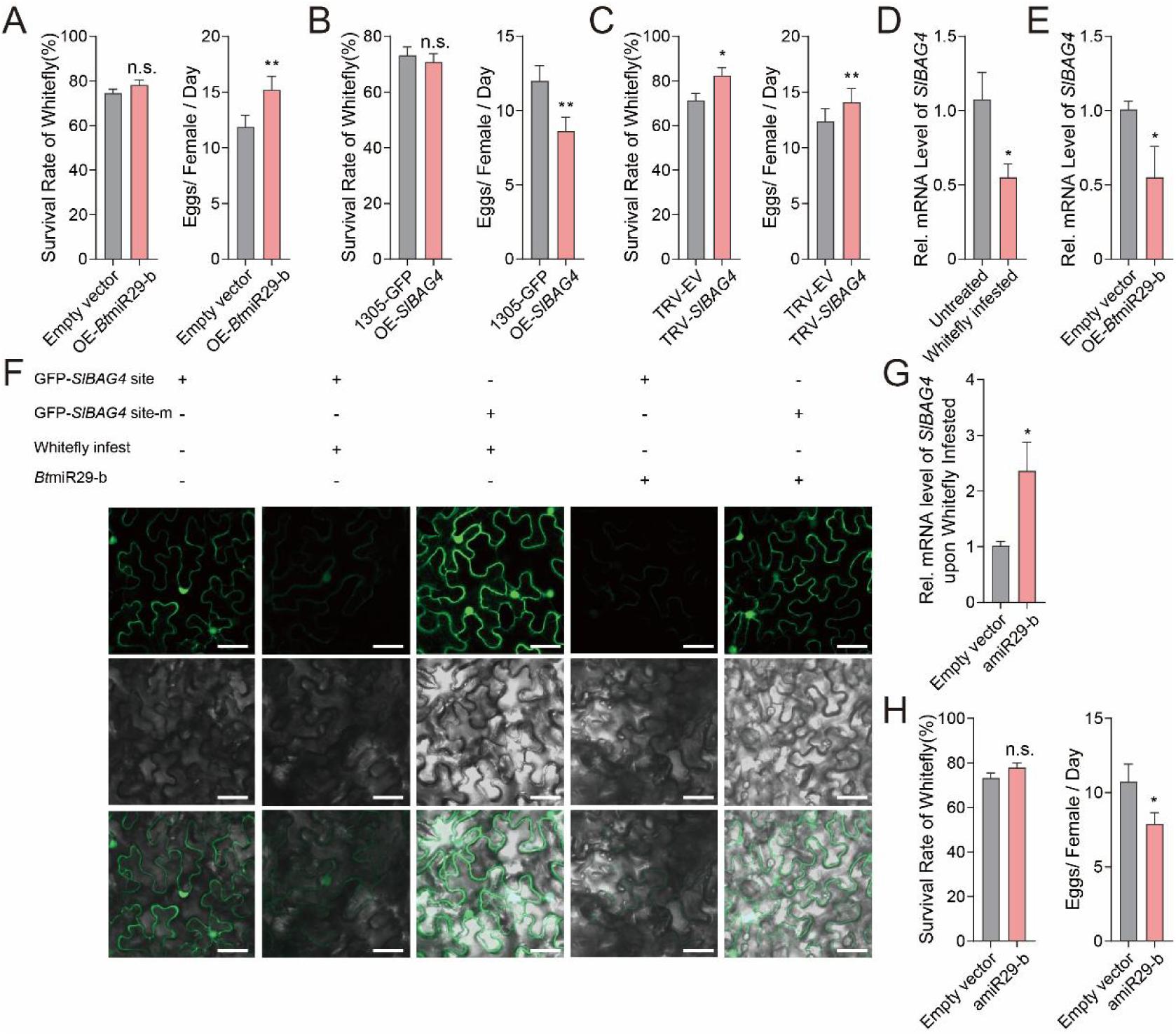
miR29-b targets defense gene *SlBAG4* in tomato. (**A**) Whiteflies performed better on tomato plants overexpressing *Bt*miR29-b. (**B**) Whiteflies perform worse on *SlBAG4*-overexpressing tomatoes. (**C**) Whiteflies perform better on *SlBAG4*-silenced tomatoes. (**D**) Whitefly infestation suppresses *SlBAG4* gene expression in tomatoes. (**E**) Overexpression of *Bt*miR29-b suppresses *SlBAG4* gene expression in tomatoes. (**F**) GFP-sensors demonstrate suppression in tomato plants upon whitefly infestation and *Bt*miR29-b overexpression. Scale bars, 40 μm. (**G**) amiR29-b overexpression in tomatoes increases *SlBAG4* mRNA levels upon whitefly feeding. (**H**) amiR29b overexpression enhances tomato defense against whitefly. Values are mean ± SEM; *n* = 40 for **A** and **H**; *n* = 25 for **B** and **C**; *n* = 6 for **D**, **E**, and **G**. Student’s *t*-test (two-tailed) was used for significant difference analysis. n.s., not significant; *, *P* < 0.05; **, *P* < 0.01.

## Discussion

Previously reported effectors of herbivore insects include proteins (*1*, *3*, *18*), chemicals, and insect transcripts (long non-coding RNA) (*38*), while no verified sRNA effector was recorded before. Although whitefly miRNAs can be detected in the phloem sap of whitefly-infested tomato plants, including *Bt*miR29-b (*21*), their role as insect effectors has not been verified. Here, we employed a series of measures from both entomological and botanical studies and confirmed the function of insect miRNA as a salivary effector **(Fig. 9)**. Our work provides new insights into the regulation of plant defense-related post-transcriptional processes by herbivore insect effectors.

**Fig. 9.**
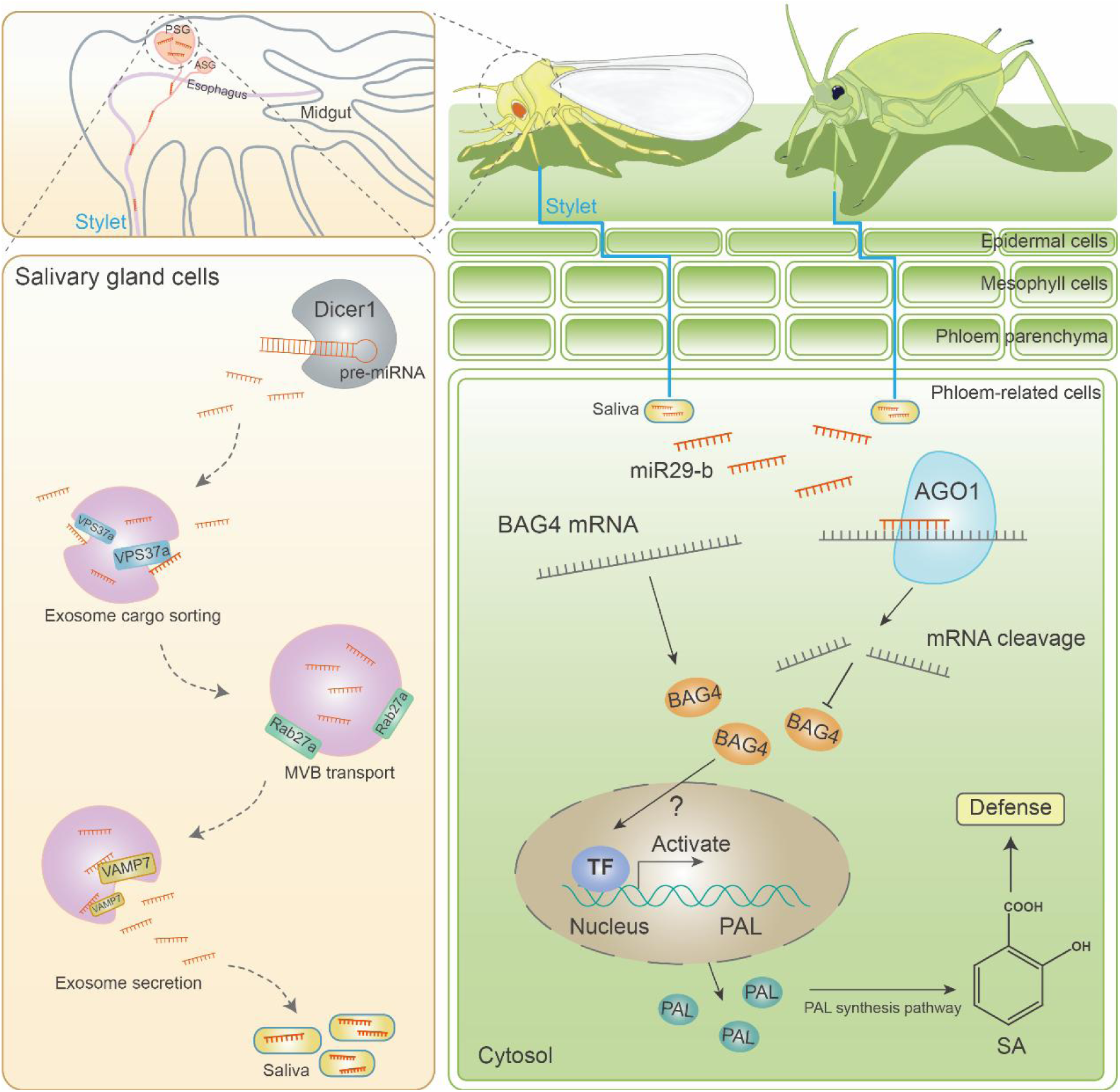
Schematic overview of herbivore insect miR29-b effector suppressing plant defense by cross-kingdom gene silencing. MiR29-b originates from the whitefly salivary gland (PSG: Primary salivary gland; ASG: Accessary salivary gland) and is transmitted to plants. In salivary gland cells, *Bt*Dicer1 produces mature miRNA and exosomes are involved in transferring and releasing it into saliva (left panel). Both whiteflies and aphids can release miR29-b into the phloem of the host plant by injecting saliva. MiR29-b hijacks the host plant AGO1 and silences the host defense gene *BAG4*. BAG4 activates the expression of the SA synthesis gene *PAL* through an unknown mechanism, thereby promoting SA accumulation (right panel). Overall, phloem-feeding insects interfere with plant SA-related defense by inhibiting BAG-mediated SA accumulation through an sRNA effector. (Abbreviation: MVB, multivesicular bodies; PAL, phenylalanine ammonia-lyase; TF, transcription factor)

We found that insect effectors *Bt*miR29-b and *Mp*miR29-b were abundant in insect salivary glands and secreted saliva **(Fig. 1C, and Fig. 7A)**. In addition, we found that *BtDcier1* but not *BtDcier2* is involved in miRNA effector formation **(Fig. 3, E to J and fig. S9)**, and *BtDcier1* is highly expressed in the salivary gland **(Fig. 3D)**. In contrast, both *BcDicer-like 1* and *BcDicer-like 2* showed functional redundancy in fungal pathogens in *Botrytis cinerea* (*39*). Moreover, we determined that exosomes play a critical role in transferring sRNA effectors from the herbivore insect whitefly to their hosts **(Fig. 4)**, expanding the knowledge of insect exosome function. Several important factors in this process were identified **(Fig. 4 and fig. S10)** and some of these factors are also highly expressed in the whitefly salivary gland **(Fig. 4, B and F)**, especially *BtVAMP7* **(Fig. 4F)**, which could release sRNA effectors carried by exosomes from the whitefly to the plant. Our work revealed the entire process of the formation and transportation of insect sRNA effectors and underscored the importance of insect salivary glands in overcoming plant resistance **(Fig. 9, left panel)**. Notably, these key factors involved in herbivore insect miRNA effector activity could be used as targets for future pest control. By silencing *BtDcier1* and exosome-related genes in whitefly, less miRNA effector was delivered into the plant and the level of *NtBAG4* and *NtPAL2* gene was much higher in tobacco upon whitefly infestation, therefore greatly reduced whitefly performance **(Fig. 3**; **Fig. 4; fig. S9; fig. S10)**.

Feeding behaviors have shaped herbivore strategies to evade host defense, particularly for phloem-feeding herbivores that coexist with their host for extended periods. Phloem-feeding herbivores extract nutrition from the plant’s phloem and release effectors into it (*15*). Previous studies have shown that plant sRNAs can transfer in the phloem, and AGO1 protein can bind sRNAs in plant phloem (*12*). In this study, we found that *Bt*miR29-b also hijacks host AGO1 and targets host genes in the phloem **(fig. S8C)**, thus enabling phloem-feeding herbivores to rapidly and continuously repress host defense. Our work highlights the pivotal role of phloem in the interaction between plants and phloem-feeding insects **(Fig. 9)**, reminding us to focus on and strengthen phloem-related crop pest resistance in the future.

It is increasingly evident that interfering with the phytohormone pathway is an efficient strategy for insects to fine-tune plant defense (*1*, *2*, *15*). SA is a well-known plant defense signal that mediates response and defense against phloem-feeding insects. The protein effector of whitefly, Armet, can decrease SA levels and the expression of SA-sentinel genes in plants (*3*). Our study shows that whiteflies can also apply this strategy through sRNA effectors. We found that BAG4, one of the targets of *Bt*miR29b, positively regulates SA levels in tobacco **(Fig. 6A)**. Similarly, in rice, the OsBAG4 protein promotes immunity and antimicrobial defense by activating SA (*40*), while we further demonstrate that BAG4 promotes SA by inducing PAL pathways **(Fig. 6, B and C)**. *Bt*miR29b suppressed the mRNA level of *NtPAL2* through silencing *NtBAG4*, which may impact PAL-mediated SA accumulation **(Fig. 5, A to D; Fig. 6, E to H)**. More studies can focus on the specific mechanism of BAG4-mediated *PAL* gene transcriptional activation **(Fig. 9, right panel)**. In addition, the activation of the PAL pathway will also promote the production of other anti-insect metabolites besides SA, such as lignin and flavonoids (*41*), which will also broaden our understanding of the BAG4-mediated herbivore-stress response. Furthermore, whitefly effectors conquer plant defense with different modes of action besides suppressing SA (*15*), such as impacting the downstream jasmonate (JA)-mediated defenses (*42*) and interfering with the interaction between WRKY33 and a central regulator in the mitogen-activated protein kinase (MAPK) cascade (*18*). In this study, we also predicted other possible target genes of *Bt*miR29-b **(Table S1)**, indicating that *Bt*miR29-b may regulate multiple plant defense mechanisms by cleaving different genes.

MiR29-b is conserved in Hemiptera insect species **(fig. S11A)**, and its target *BAG4* also exists in a wide range of angiosperm species **(fig. S12)**. Despite the different target sites, the detection of the expression of GFP fused with the target sites, and gene expression tests revealed that insect sRNA effectors can cleave *BAG4* genes in different Solanaceae plants **(Fig. 5**; **Fig. 8)**, suggesting that phloem-feeding insects may overcome the conserved defense genes of different host plants using a common effector. These results are a good illustration of the arms race between phloem-feeding insects and their host plants, with both sides gaming around conservative SA-mediated defense **(Fig. 6)**, as a result of lengthy co-evolution. Significantly, our research demonstrates that the cohort of phloem-feeding insects shares common strategies for suppressing host defense mechanisms. Moreover, their chosen targets exhibit widespread prevalence across various plant species. This revelation underscores the importance of directing our focus toward these conserved mechanisms in enduring plant-insect interaction. By doing so, we can simultaneously mitigate the damage caused by these insect pests to crops. For example, within the scope of this study, the amplification of amiRNA was found to bolster the plant’s defensive capabilities against both whiteflies and aphids **(Fig. 2C**; **Fig. 7D**; **Fig. 8H)**.

Overall, our findings imply that the sRNA of phloem-feeding insects acts as the effector to fine-tune plant defense, and cross-kingdom sRNA-mediated gene silencing may be widespread in the interactions between phloem-feeding insects and plant species. This insight deepens our understanding of insect effectors and their target genes, as well as the mechanisms underlying their functions. Meanwhile, based on the investigation of natural cross-kingdom gene transfer between insects and plants, some pest control measures can be proposed. The conserved anti-whitefly gene *BAG4* can be a candidate goal for transgenic crops, and sRNA-related insect genes can be terrific targets for the promising RNA interference (RNAi)-based insect pest control strategy.

## Materials and methods

### Plants and insects

MEAM1 *B. tabaci* were reared on healthy cotton (*Gossypium hirsutum* cv. ZheMian 1793), and *M. persicae* were reared on healthy tobacco plants (*N. tabacum* cv. NC89) in climate chambers at 26 ± 1℃, 60 ± 10% relative humidity, and a photoperiod of 14:10 h (L:D). Additionally, *N. benthamiana*, *Solanum lycopersicum* (tomato) cultivar Hezuo903, *A. thaliana* ecotype Col-0, and knockout mutants *ago1* (SALK_210111C) and *bag4* (SALK_027577C) were also used. The plants were individually grown in 10 cm-diameter plastic pots under the same conditions as the insects.

### Whitefly Small RNA sequencing

Total RNAs were extracted from whitefly samples using TRIzol (Invitrogen, Carlsbad, CA) according to the manufacturer’s instructions. Small RNA sequencing was performed as previously described (*22*). Purified RNA fragments were ligated to two adaptors (5’ adaptor: GTTCAGAGTTCTACAGTCCGACGATC; 3’ adaptor: TGGAATTCTCGGGTGCCAAGG) and converted to complementary DNA (cDNA) for RT-PCR amplification. Then these small RNAs were amplified with PCR, followed by sequencing with an Illumina HiSeq 2500 (Illumina, San Diego, IL, USA) to construct a library.

### Herbivore insect salivary gland and saliva collection

The adults of whiteflies were anesthetized on ice and their primary salivary glands (PSG) were dissected in 1× PBS. Then the salivary gland samples were transferred to TRIzol (Invitrogen, Carlsbad, CA) for further total RNA extraction. To collect saliva, 600 whiteflies were fed on an artificial diet (15% sucrose solution) for 8 days and saliva was collected every 24 hours. Finally, the collected saliva solution was concentrated to extract total RNAs.

### Plant phloem sap collection

Plant leaves were completely transverse with a clean scissor, and the phloem sap collections were performed as previously described (*43*). Total RNA was isolated using the Applied Biosystems Arcturus™ CapSure™ LCM MicroCaps (Thermo Fisher, Carlsbad, CA), and reverse transcription of miRNAs was performed using the miRNA 1st strand cDNA synthesis kit (Accurate Biology, Hunan, China). Subsequently, RT-PCR was used to detect miR29-b in the plant phloem sap.

### Cloning and sequencing of herbivore insect sRNA

Whiteflies and aphids were collected to verify the sequence of miR29-b. Total RNAs were isolated from the insect samples using TRIzol reagent following the instructions. The miRNA 1st strand cDNA synthesis kit (Accurate Biology, Hunan, China) was used to prepare the polyA-enriched cDNA. All PCR amplifications were carried out using 2 × SYBR Green *Pro Taq* HS Premix (Accurate Biology, Hunan, China), and amplified fragments were purified using an AxyPrepTM DNA Gel Extraction Kit (Axygen, West

Orange, NJ, USA), then cloned into the pClone007 Blunt Vector (Tsingke, Beijing, China) and sequenced. Primers used in this study are listed in **Table S3**.

### Quantitative real-time PCR analysis

Quantitative PCR of the target mRNAs and miRNAs was carried out on a CFX96 real-time PCR detection system (Bio-Rad) using SYBR Green Premix *Pro Taq* HS qPCR kit (Accurate Biology, Hunan, China) following the instructions. Expression of the *GAPDH*, *Actin*, *EF1⍺, β-tub*, *miR-275*, and *U6* snRNA genes were used as the internal controls for data analysis. The mRNA and miRNA expression levels were calculated by the 2^-ΔΔCt^ method (*44*). Primers used in this study are listed in **Table S3**.

### Insect assay

A special micro-leaf cage was set up on the test plants, and 5 male and 5 female whiteflies were released into the cage. The survival rate and egg production of whiteflies were measured 3 days later. The whiteflies in the double tube also recorded mortality after three days and the number of eggs laid on the plastic film. For aphid bioassay, three female aphids were placed in a leaf cage fixed on the leaves of the tested plants, and the daily production was counted for three consecutive days.

### GFP-reporter detection

The tested gene sequences were cloned into the pCAMBIA1305-GFP vector. Primers used in this study are listed in **Table S3**. *Agrobacterium* cultures expressing Green fluorescent protein (GFP)-tagged protein were infiltrated into the leaves of approximately 4-week-old *N. benthamiana* plants. After 48 hours of infiltration, 500 whitefly adults were placed at the infiltration site for 48 hours, then GFP fluorescence was detected using the confocal microscopy system (Zeiss LSM710, Oberkochen, Germany). GFP-tagged protein and *Bt*miR29-b were co-infiltrated and the GFP fluorescence was detected after 4 days.

### GUS-reporter detection

To construct the GUS sensors, promoters or gene sites were cloned into the pCAMBIA121-GUS vector to drive the *GUS*-reporter gene. *Agrobacterium* cultures containing the *GUS*-reporter gene were infiltrated into *N. benthamiana* or *N. tabacum* leaves. The leaf tissues were collected after 4 days of infiltration. The feeding treatment and sampling of whiteflies were the same as above. GUS detection was performed using a GUS staining detection kit (Huayueyang, Beijing, China). The leaves were incubated in the staining buffer overnight at 37°C. After staining, the leaves were cleared with 70% ethanol until the negative control turned white. The promoter of *AtSUC2* (sucrose transporters 2) served as a phloem-expression marker. Primers used in this study are listed in **Table S3**.

### Phytohormone detection

The tobacco leaves were wrapped in tin foil and immediately powdered in liquid nitrogen. Phytohormone extraction was performed by adding 1 mL of HPLC-grade ethyl acetate (Sinopharm Chemical Reagent Co., Ltd, Shanghai, China), which contained 10 ng of D4-SA, to 0.15 g of leaf powder. The samples were then vortexed for 15 minutes and centrifuged at 13000 rpm for 20 minutes at 4°C. After centrifugation, the supernatants were collected and evaporated by a vacuum concentrator (Eppendorf, Hamburg, Germany) at 30°C for about 30 minutes until dry. The residues were resuspended in 110 µL of MeOH: H_2_O (50:50, v/v) and centrifuged at 13000 rpm for 10 minutes at 4°C. The resulting supernatants were collected and analyzed using a high-performance liquid chromatography-tandem mass spectrometry system (TripleTOF 5600+; AB Sciex, Redwood City, CA, USA). Each treatment was replicated six times.

### Degradome sequencing

Degradome sequencing analysis, following the previous study (*22*), was commissioned to Lianchuan Biotechnology Co., Ltd. to conduct.

### Virus-induced gene silencing and transient gene expression

For the VIGS assay, a 250-bp fragment of *NtBAG4* (XM_016607951.1) and a conserved 400-bp fragment of four *NtAGO1* genes (LOC107806342: XM_016630482.1, XM_016630480.1, and XM_016630483.1; LOC107790785: XM_016612746.1, XM_016612745.1, and XM_016612747.1; LOC107771391: XM_016590746.1; LOC107789474: XM_016611294.1) were amplified and cloned into the pBIN2mDNA1 vector, and a 300-bp fragment of *SlBAG4* (XM_004240364.4) was amplified and cloned into the pTRV-RNA2 vector. For the transient expression assay, the full length of *NtBAG4* and *SlBAG4* were cloned into the plant expression vector pCAMBIA1305. All of these constructed vectors were transformed into *A. tumefaciens* strain EHA105 using electroporation. The *A. tumefaciens* cells were grown at 28°C and 200 rpm until the OD_600_ reached approximately 1.0. Then, the cultures were collected and resuspended in infiltration buffer (100 mM 2-(N-morpholino) ethanesulfonic acid (MES), 10 mM MgCl_2_, 2 mM acetosyringone) at room temperature for 3 hours. Finally, the suspensions were used to infiltrate plant leaves using 1 ml blunt syringes. Primers used in this study are listed in **Table S3**.

### Bioinformatics analysis

Multiple sequence alignments were performed by ClustalW, and the phylogenetic trees were constructed using the maximum likelihood method based on the Whelan Goldman (WAG) model with 1000 bootstrap replications in MEGA7.0 software (*45*). The sequences used for phylogenetic analysis are listed in **Table S4**.

### Production of dsRNA transcription templates and synthesis of dsRNA

To identify the genes involved in the exosome system of whiteflies, we conducted BLASTP and TBLASTN searches on the MEAM1 *B. tabaci* genome using established key genes of insect exosome systems as queries (*46*). These key genes include *CD63*, *Syntenin*, *SMPD* (involved in exosome formation), *VPS37a* (involved in cargo sorting), *Rab27a* (involved in the transport process), and *VAMP7* (involved in exosome secretion) (*47*). To generate double-stranded RNA (dsRNA), we amplified fragment templates of *Bt-Dicer 1* (Bta12886), *Bt-Dicer 2* (Bta10685), and the aforementioned key genes of the exosome system (CD63: Bta11700; syntenin: Bta13200; SMPD: Bta11772; VPS37a: Bta11125; Rab27a: Bta08735; VAMP7: Bta10299) using PCR with primers that contained the *T7* promoter sequence at the 5’ ends **(Table S3)**. The dsRNAs utilized in this study were synthesized employing the T7 High Yield RNA Transcription Kit (Vazyme, Nanjing, China) following the provided instructions.

Gene silencing of the target genes in whiteflies was performed using the parafilm clip nutrient solution method as previously described (*48*). The dsRNA was diluted into 15% (w/v) sucrose solution at a concentration of 200 ng/μL. Approximately 100 whitefly adults were collected and placed into glass tubes (1.5 cm in diameter and 10 cm in height). After a 24-hour feeding, the RNAi efficiency was analyzed by qPCR. Bioassays of gene-silenced whiteflies on tobacco were performed as described above.

### MicroRNA agomir and antagomir treatment in *vivo*

Agomir and antagomir (*49*) were used to study the function of miR29-b in *vivo*. Agomir29-b and antagomir29-b were diluted into a 15% (w/v) sucrose solution at a concentration of 100 ng/μL. Feeding methods are the same as described above. Primers used in this study are listed in **Table S3**.

### Vector construction of artificial miRNA and artificial target mimic

Artificial miRNA (amiRNA) is a recently developed method for overexpressing miRNA with high specificity, while target mimicry is a natural mechanism that inhibits miRNA function. To study the function of miRNAs, we constructed amiRNA and artificial mimics. The precursor of amiRNA was generated following the method previously described (*50*), and the artificial mimic was generated following the previous method (*23*). The precursor sequences of amiRNA and mimic were then synthesized by GenScript (Nanjing, China). Finally, the precursors were inserted into the pC1300-CaMV 35S-nos vector between the *CaMV 35S* promoter and the nos terminator to form the final expression vectors. Primers used in this study are listed in **Table S3**.

### Statistical analysis

One-way ANOVA followed by Fisher’s least significant difference test and Student’s *t*-tests using SPSS 26.0 were carried out in this work and are shown in the figure legends.

## Acknowledgments

We thank Rong Jin from Agricultural Experiment Station, Zhejiang University for greenhouse management.

## Funding

Financial support for this study was provided by: National Natural Science Foundation of China (31925033, 32161143008) National Key Research and Development Program (2021YFC2600100).

## Author contributions

Conceptualization: W.H. and X.W.

Methodology: W.H., S.J., F.Z., H.S., J.W., and R.X.

Investigation: W.H. and X.W.

Visualization: W.H. and S.J.

Supervision: W.H., S.J., and X.W.

Writing—original draft: W.H. and S.J.

Writing—review & editing: W.H., F.Z., and X.W.

## Competing interests

Authors declare that they have no competing interests.

## Data and materials availability

All data needed to evaluate the conclusions in the paper are present in the paper and/or the Supplementary Materials.

## Supplementary Materials

**This PDF file includes**:

Supplementary Text

Figs. S1 to S13

**Other Supplementary Material for this manuscript includes the following**:

Tables S1 to S4

## Notes

### Competing Interest Statement

The authors have declared no competing interest.

