## Supplementary Figures for "Herbivore insect small RNA effector suppress plant defense by cross-kingdom gene silencing"

Wen-Hao Han *et al.*

**This PDF file includes:**

Supplementary Text
Figs. S1 to S13

**Other Supplementary Materials for this manuscript include the following:**
Tables S1 to S4

**Supplementary Text**

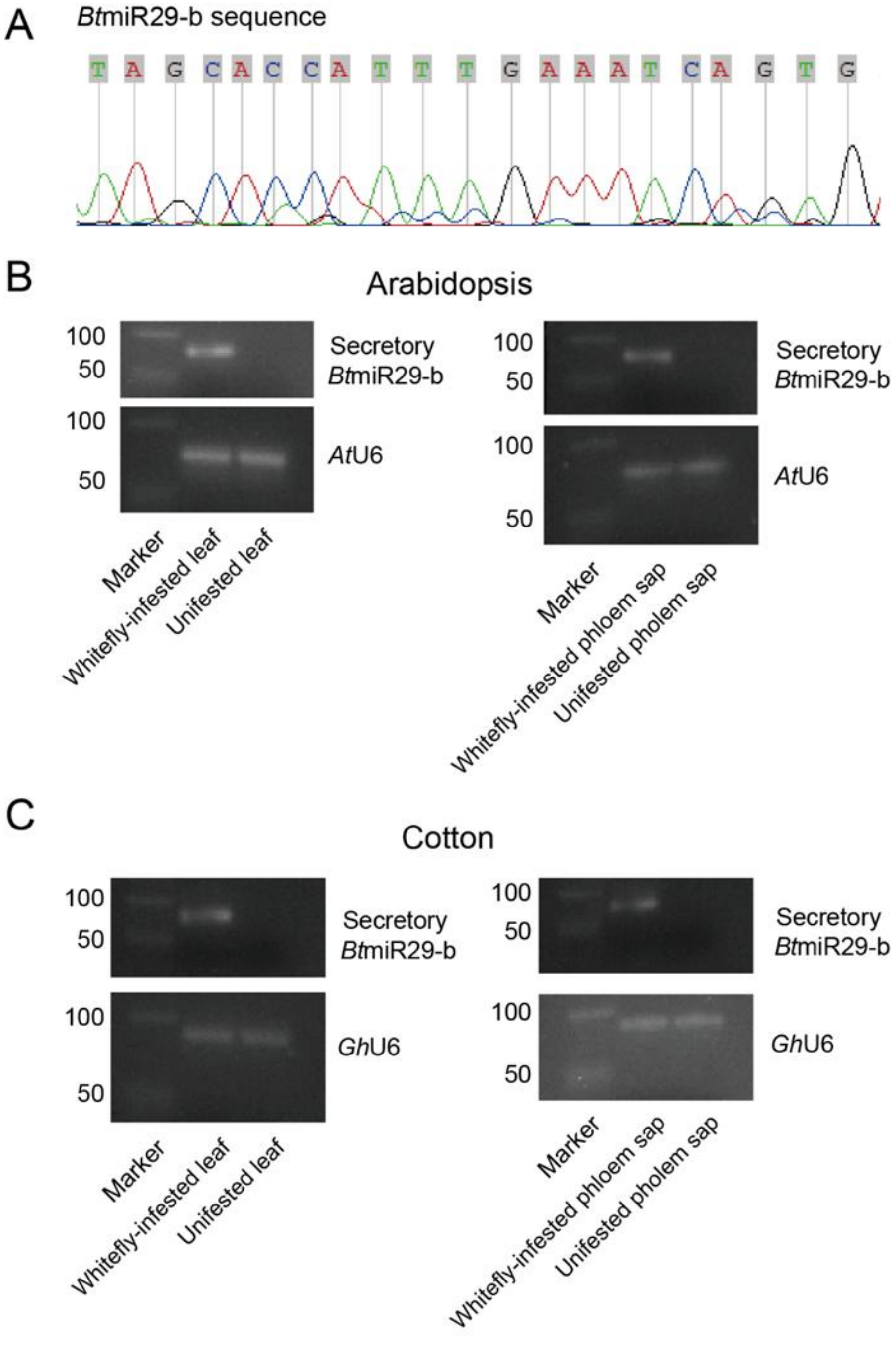


**Fig. S1.**

(A) The sequence of *Bt*miR29-b was determined through sequencing. (B) *Bt*miR29-b was detected in the whitefly-infested Arabidopsis host and its phloem sap. (C) *Bt*miR29-b was detected in whitefly-infested cotton and its phloem sap.

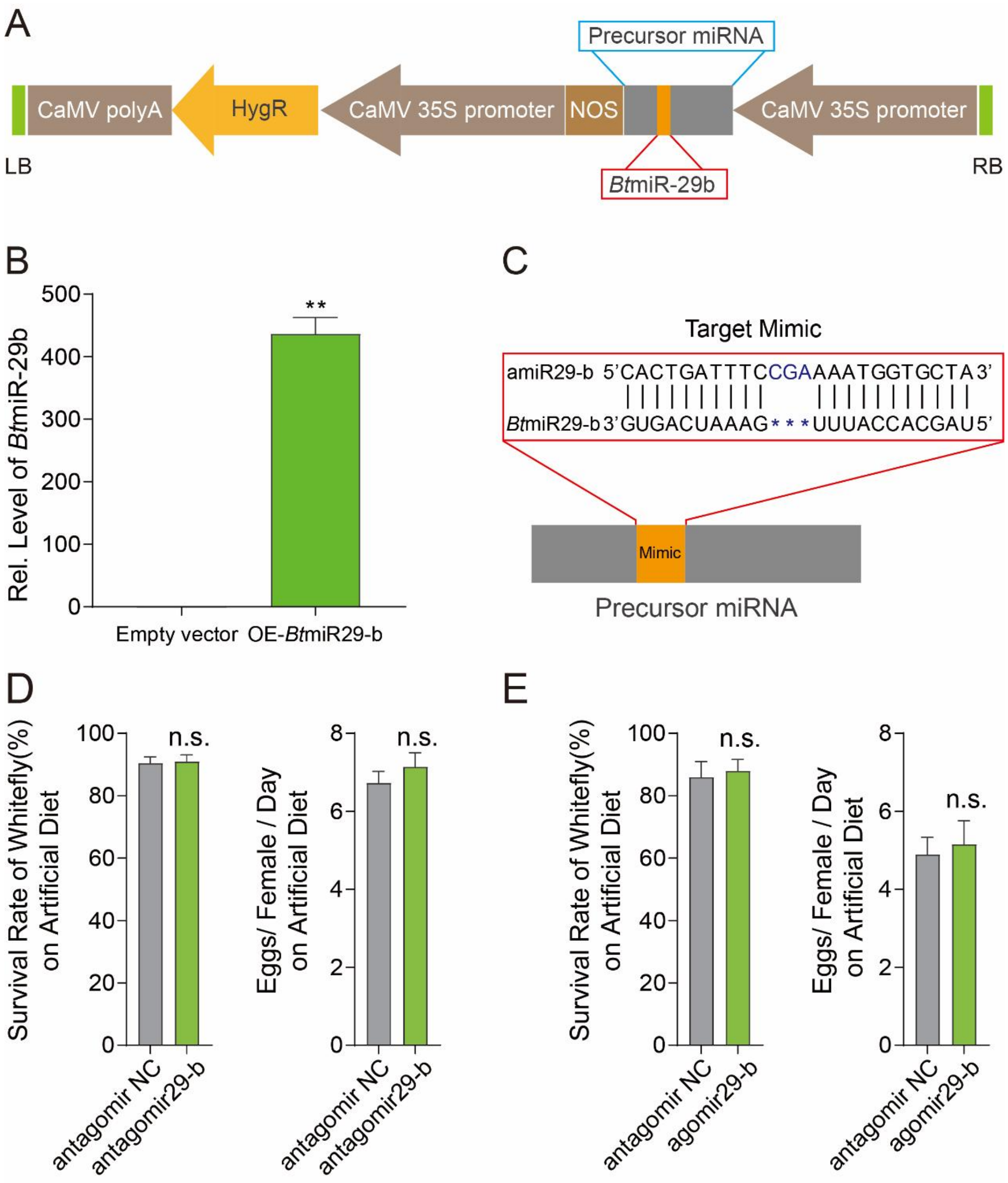


**Fig. S2.**

(A) Schematic diagram of the *Bt*miR29-b expression cassette *p35S-Bt*miR29-b overexpressed in tobacco plants through *Agrobacterium*-mediated transformation. The blue box represents the miRNA precursor, and the red box represents *Bt*miR29-b. HygR: Hygromycin B phosphotransferase; NOS: Nos Terminator; RB: Right Border; LB: Left Border. (B) Relative mRNA level of *Bt*miR29-b significantly increased in *Bt*miR29-b-overexpressed tobacco plants. (C) Design of mimicry sequences (amiR29-b) for *Bt*miR29-b. (D) Three-day survival and fecundity of whiteflies feeding with antagomir29-b were unaffected under suitable conditions provided with artificial feed. (E) The performance of whiteflies feeding with agomir29-b was unaffected under suitable conditions provided with an artificial diet. Values are mean ± SEM; $n$ = 6 for B; $n$ = 20 for D and E. Student's $t$-test (two-tailed) was used for significant difference analysis. n.s., not significant; **, $P < 0.01$.

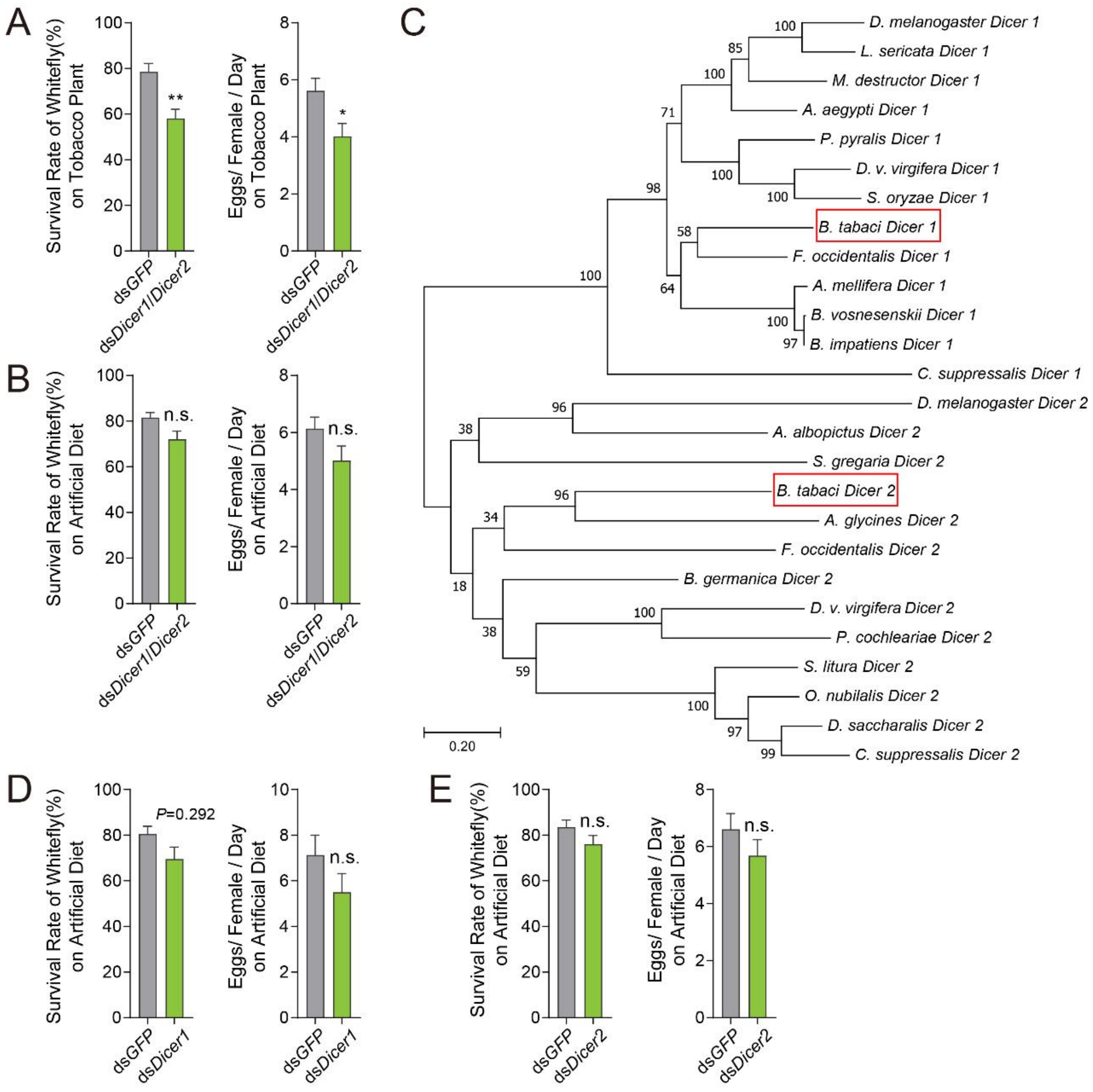


**Fig. S3.**

(A) *BtDicer1*/*Dicer2*-silenced whiteflies perform worse on tobacco host plants. (B) The performance of whiteflies with *BtDicer1*/*Dicer2* silencing was unaffected under suitable conditions provided with an artificial diet. (C) Phylogenetic tree of Dicer proteins in insect species. (D) The performance of *BtDicer1*-silenced whiteflies under suitable conditions provided with an artificial diet was unaffected. (E) The performance of *BtDicer2*-silenced whiteflies under suitable conditions provided with an artificial diet was unaffected. Values are mean ± SEM; $n$ = 20 for A, B, D, and E. Student's $t$-test (two-tailed) was used for significant difference analysis. n.s., not significant; *, $P < 0.05$; **, $P < 0.01$.

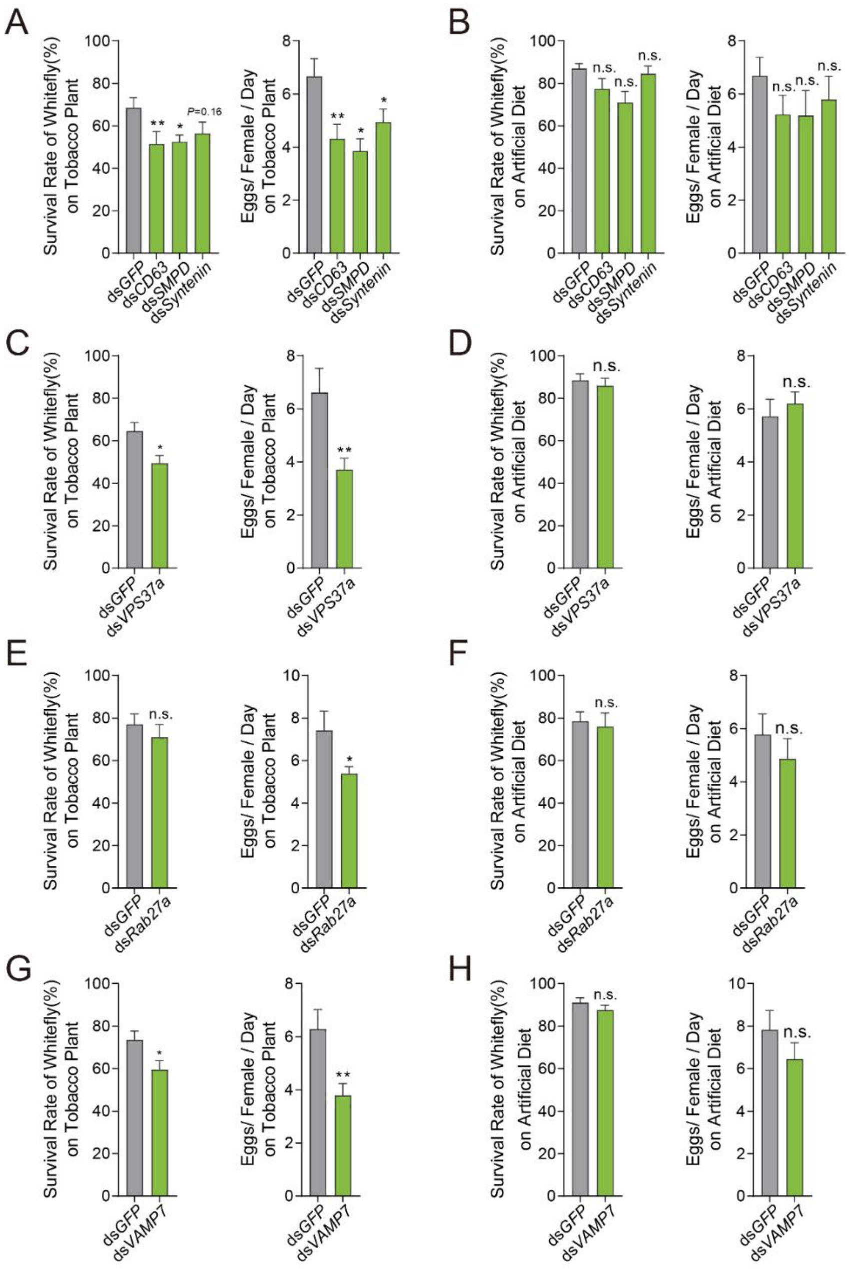


**Fig. S4.**

(A) The performance of *BtCD63*, *BtSMPD*, and *BtSyntenin*-silenced whiteflies was decreased on tobacco host plants. (B) The performance of *BtCD63, BtSMPD*, and *BtSyntenin*-silenced whiteflies was not affected under suitable conditions provided with an artificial diet. (C, D) *BtVPS37a*-silenced whiteflies perform similarly on an artificial diet but worse on tobacco host plants. (E, F) *BtRab27a*-silenced whiteflies perform similarly on an artificial diet but worse on tobacco host plants. (G, H) *BtVAMP7*-silenced whiteflies perform similarly on an artificial diet but worse on tobacco host plants. Values are mean ± SEM; $n = 20$ for A, B, C, D, E, F, G, and H. Student's *t*-test (two-tailed) was used for significant difference analysis. n.s., not significant; *, $P < 0.05$; **, $P < 0.01$.

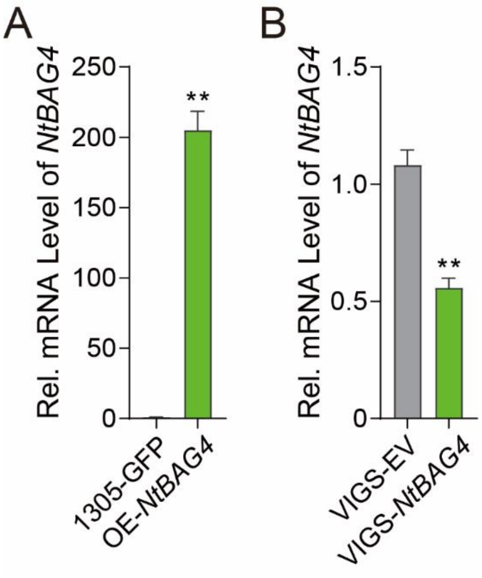


**Fig. S5.**

(A) Relative mRNA level of *NtBAG4* significantly increased in *NtBAG4*-overexpressed tobacco plants. (B) Relative mRNA level of *NtBAG4* significantly decreased in *NtBAG4*-silenced plants. Values are mean ± SEM; $n = 6$. Student's *t*-test (two-tailed) was used for significant difference analysis. **, $P < 0.01$.

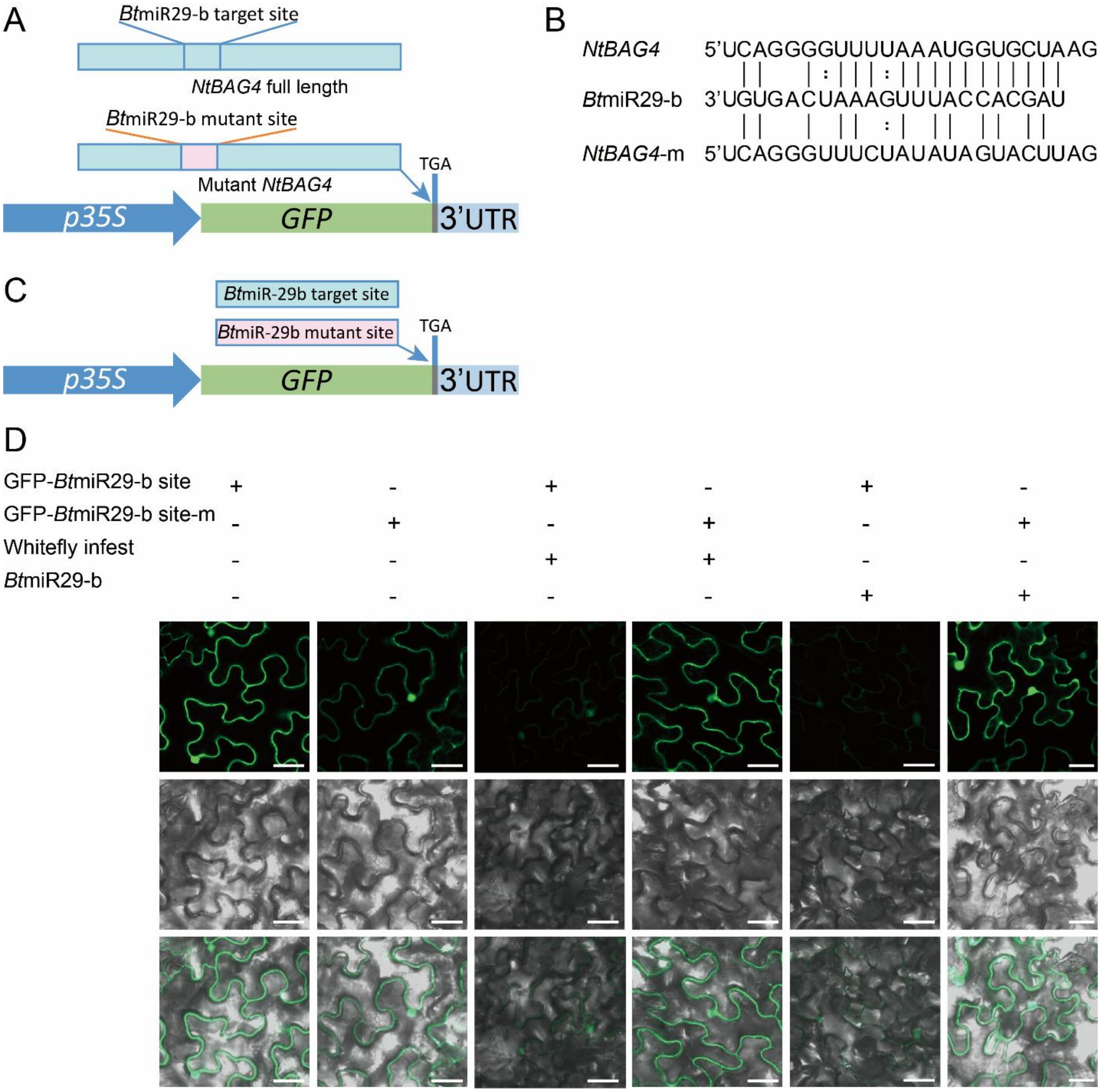


**Fig. S6.**

(A) Schematic diagram of the GFP sensor carrying *NtBAG4* and *NtBAG4* mutant site. TGA: Termination Codon; UTR: Untranslated Regions. The blue long strip represents the full length of the *NtBAG4* transcript, and the blue short strip in the full length represents the *Bt*miR29-b target site at 245-268bp. The pink strip in the blue long strip represents the mutant *Bt*miR29-b target site at the same transcript position. (B) Target site and target site mutated versions of *Bt*miR29-b for the tobacco plant target gene *NtBAG4* used in this study. (C) Schematic diagram of the GFP sensor carrying *Bt*miR29-b target site and mutant target site. TGA: Termination Codon; UTR: Untranslated Regions. (D) GFP sensors with the *Bt*miR29-b target sites of *NtBAG4* or target site-m showed differential expression during whitefly infestation. (Top) GFP. (Middle) Bright-field. (Bottom) GFP/bright-field overlay. Scale bars, 40 μm.

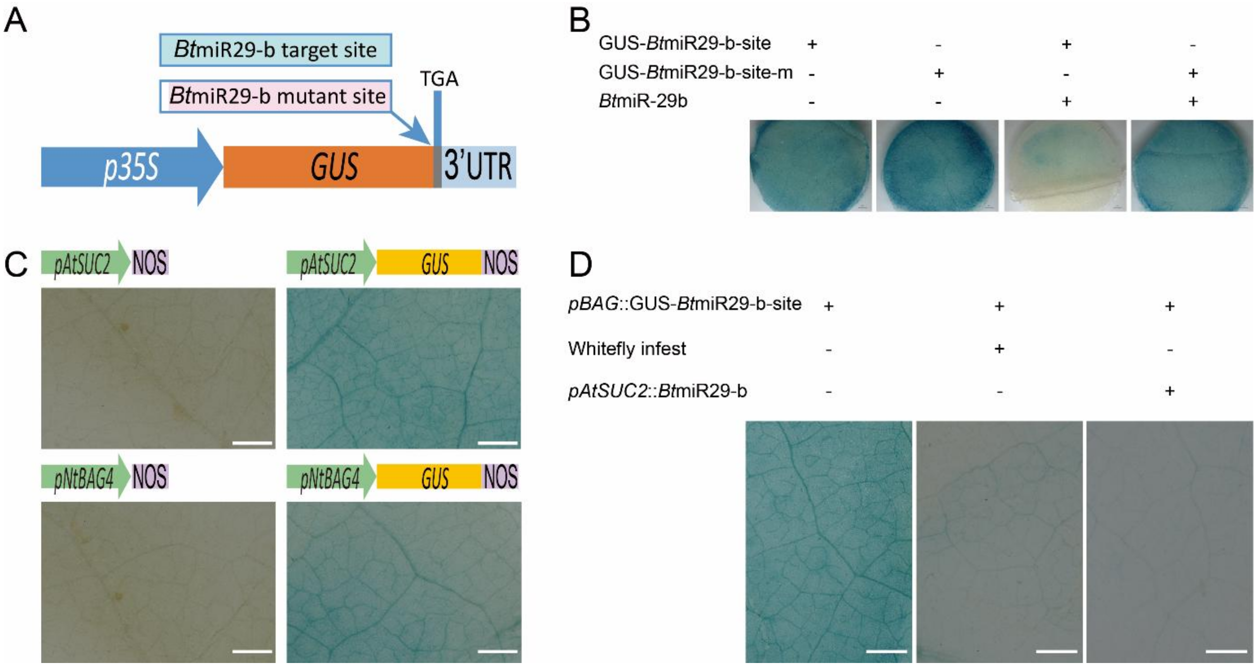


**Fig. S7.**

(A) Schematic diagram of the GUS sensor carrying the *Bt*miR29-b target site and mutant site. TGA: Termination Codon; UTR: Untranslated Regions. (B) GUS sensors confirmed *Bt*miR29-b's impact on its target site of *NtBAG4* in *Nicotiana tabacum*. (C) *NtBAG4* is expressed in the phloem of *N. benthamiana*. *pAtSUC2*: Promoter of *AtSUC2* (*Sucrose transporters*, a phloem-expression marker); *pNtBAG*: Promoter of *NtBAG4*. NOS: Nos Terminator. (D) Cleavage of the host target *NtBAG4* by whitefly or *Bt*miR29-b occurs in the host phloem of *N. benthamiana*, as shown by the GUS sensor. Scale bars, 1 mm.

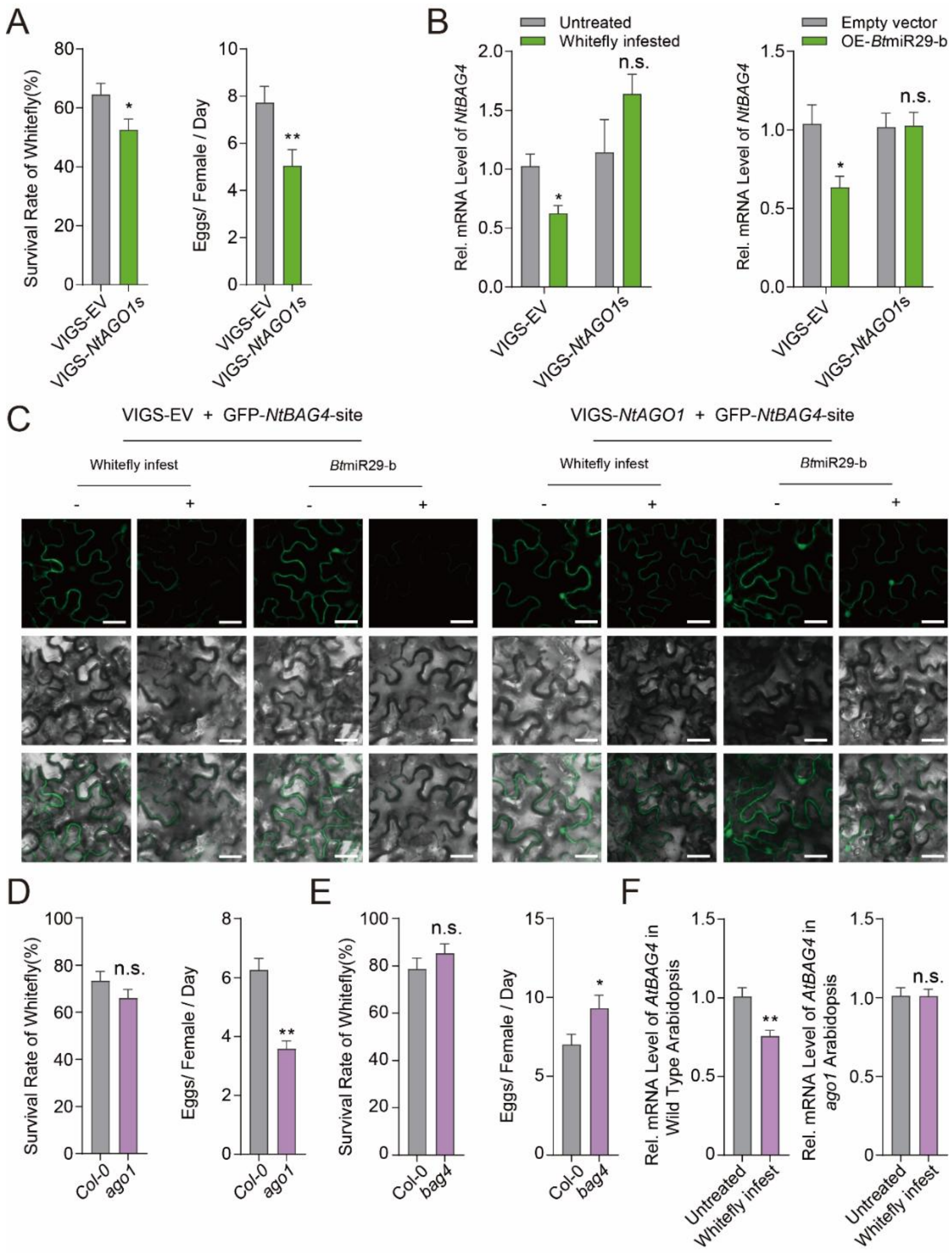


**Fig. S8.**

(A) Silencing *NtAGO1*s in host tobacco enhanced defense against whiteflies. (B) *NtAGO1*s-silenced tobacco did not inhibit the expression of *NtBAG4* upon whitefly infestation and *Bt*miR29-b overexpression. (C) GFP-sensors confirmed the suppression of host *NtBAG4* in *NtAGO1*s-silenced tobacco during whitefly infestation and *Bt*miR29-b overexpression. Scale bars, 40 μm. (D) *ago1* Arabidopsis exhibited improved defense against whiteflies. (E) *AtBAG4* played a role in Arabidopsis's defense against whiteflies. (F) Whitefly infestation suppressed *AtBAG4* in wild-type Arabidopsis, not in *ago1* plants. Values are mean ± SEM; $n = 20$ for A, $n = 6$ for B and F; $n = 15$ for D and E. Student's *t*-test (two-tailed) was used for significant difference analysis. n.s., not significant; *, $P < 0.05$; **, $P < 0.01$.

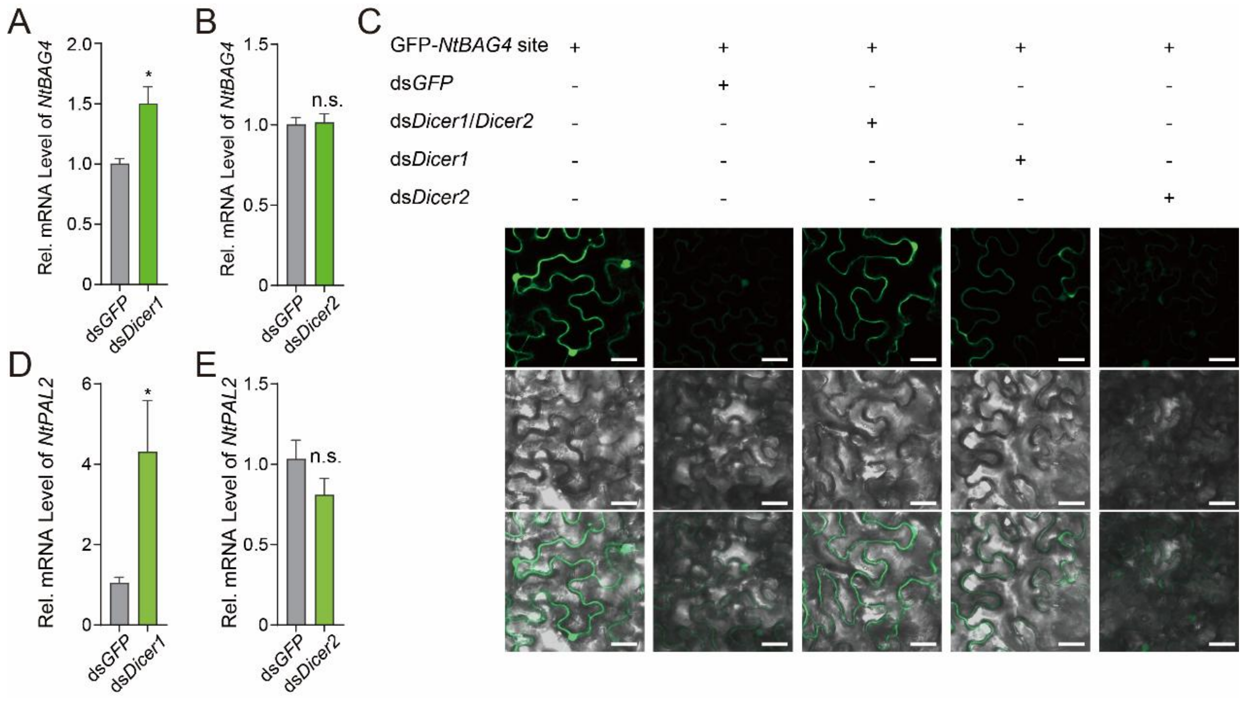


**Fig. S9.**

(A) *NtBAG4* mRNA level significantly increased in tobacco infested by *BtDicer1*-silenced whiteflies. (B) *NtBAG4* mRNA level remained unchanged between *BtDicer2*-silenced and control whiteflies. (C) GFP-*NtBAG4* site sensor silencing by ds*GFP* and ds*Dicer2* whiteflies, not by ds*Dicer1* whiteflies. Scale bar, 40 mm. (D, E) *NtPAL2* mRNA is higher in tobacco infested by *BtDicer1*-silenced whiteflies, not *BtDicer2*-silenced whiteflies. Values are mean ± SEM; $n$ = 6 for A, B, D, and E. Student's $t$-test (two-tailed) was used for significant difference analysis. n.s., not significant; *, $P < 0.05$.

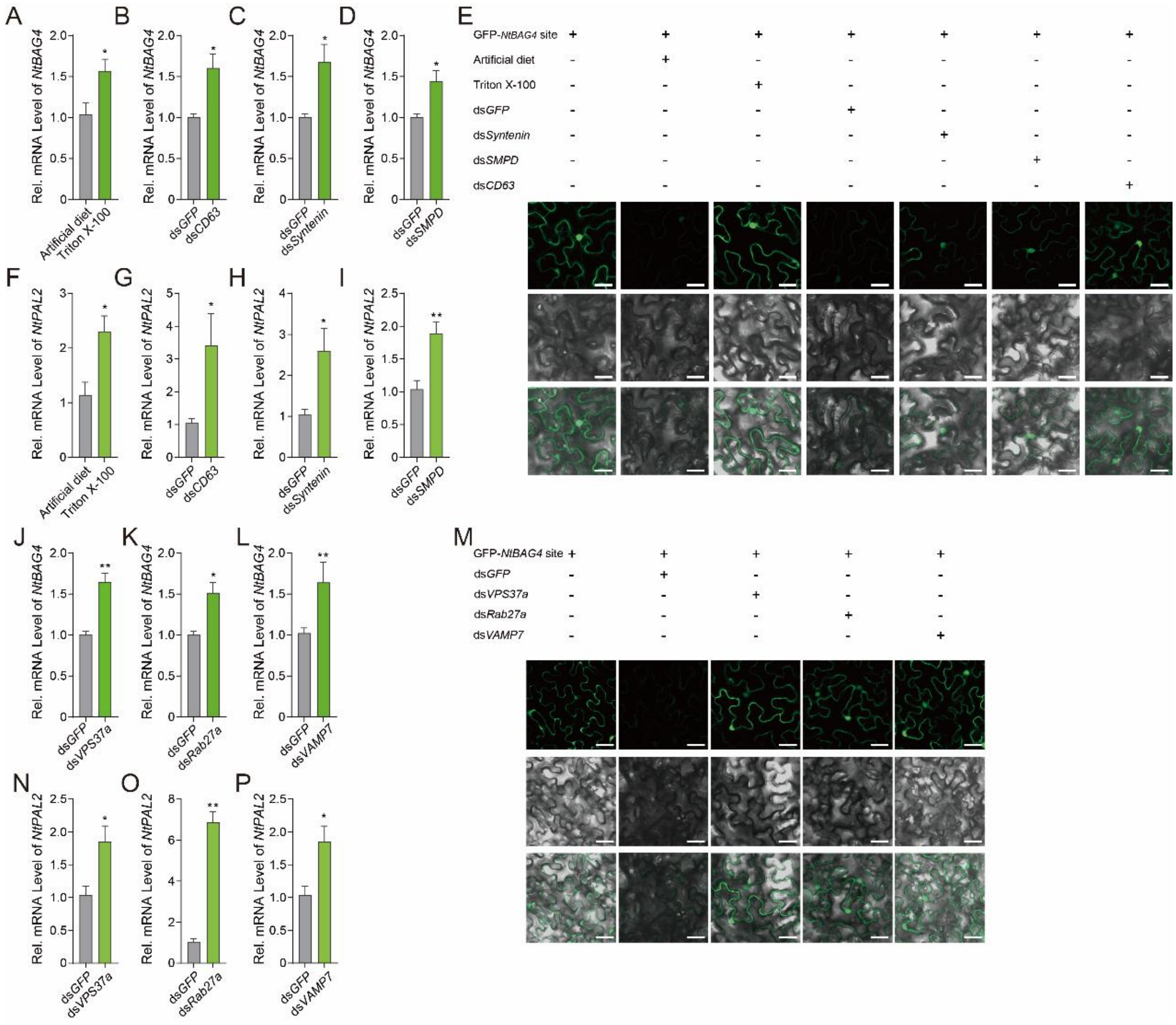


**Fig. S10.**

(A, B, C, D) Feeding 0.01 mM TritonX-100 and silencing *BtCD63*, *BtSyntenin*, and *BtSMPD* genes in whiteflies led to significantly higher *NtBAG4* mRNA levels in infested tobacco. (E) GFP-*NtBAG4* site sensor silencing by artificial diet-feeding and ds*GFP* whiteflies, not by Triton X-100-feeding, ds*Syntenin*, ds*SMPD*, and ds*CD63* whiteflies. Triton X-100 mixed with artificial diet (15 % sucrose water); the treatment of the artificial diet in the graph is a negative control without Triton X-100. (F, G, H, I) *NtPAL2* mRNA is higher in tobacco infested by whiteflies fed with Triton X-100, ds*CD63*, ds*Syntenin*, and ds*SMPD*. (J, K, L) Silencing *BtVPS37a*, *BtRab27a*, and *BtVAMP7* genes in whiteflies led to significantly higher *NtBAG4* mRNA levels in infested tobacco. (M) The GFP-*NtBAG4* site sensor was not silenced by ds*VPS37a*, *dsRab27a*, and *dsVAMP7* whiteflies. (N, O, P) *NtPAL2* mRNA is higher in tobacco infested by *BtVPS37a*, *BtRab27a*, and *BtVAMP7*-silenced whitefly. Scale bar, 40 mm. Values are mean ± SEM; $n = 6$. Student's *t*-test (two-tailed) was used for significant difference analysis. *, $P < 0.05$; **, $P < 0.01$.

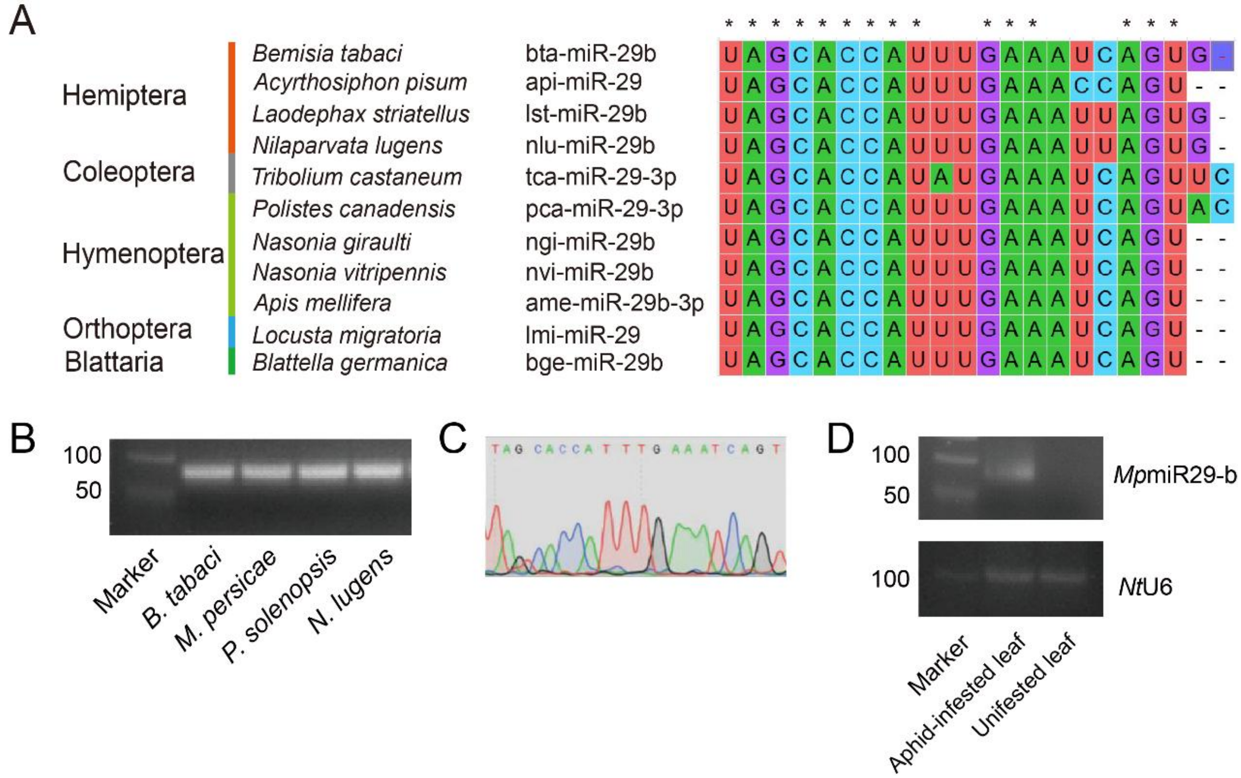


**Fig. S11.**

(A) *Bt*miR29-b is conserved across various insect species. (B) miR29-b was detected in several Hemipterous insect species. (C) The sequence of *Mp*miR29-b was confirmed by sequencing. (D) *Mp*miR29-b was detected in aphid-infested tobacco plant leaves.

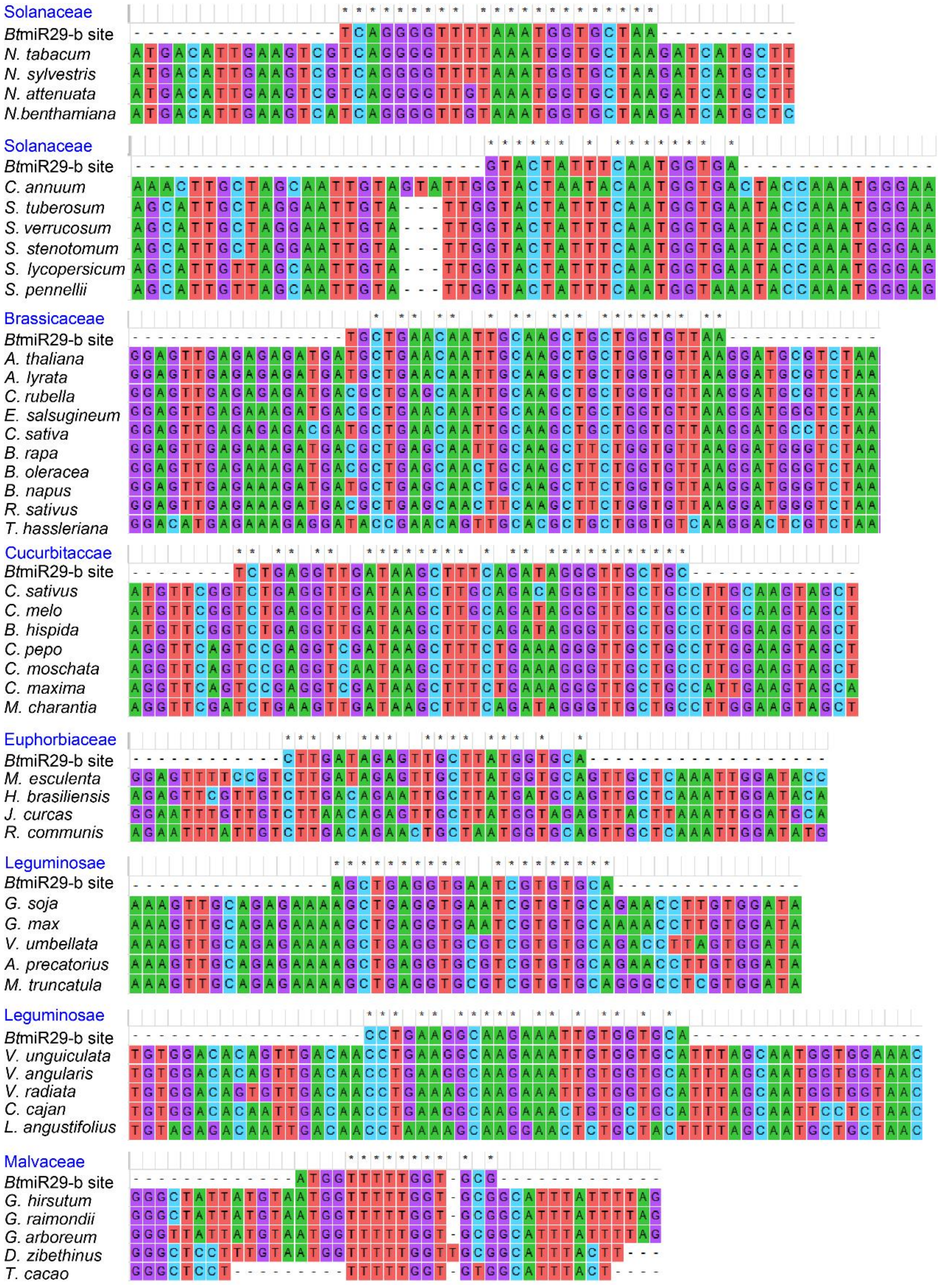


**Fig. S12.**

The cleavage site of *Bt*miR29-b in *BAG4* transcripts in different plant species including Solanaceae (*Nicotiana* plants and other species), Brassicaceae, Fabaceae, Euphorbiaceae, Cucurbitaccae, and Malvaceae plants predicated by Target Finder.

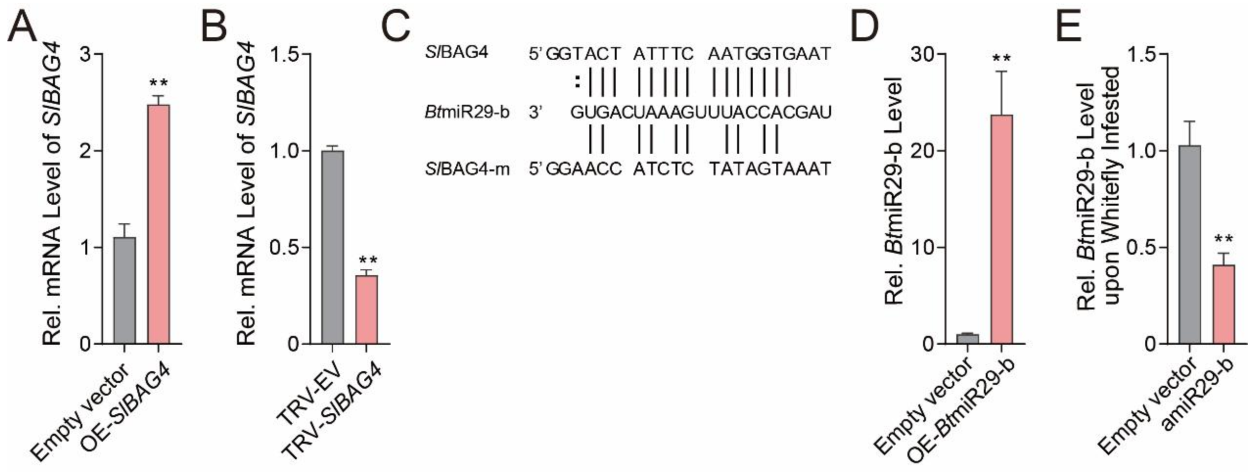


**Fig. S13.**

(A) Relative mRNA level of *SlBAG4* significantly increased in *SlBAG4*-overexpressed tomato plants. (B) Relative mRNA level of *SlBAG4* significantly decreased in *SlBAG4*-silenced plants. (C) Target site and target site mutated versions of *Bt*miR29-b for tomato target gene *SlBAG4* were used in this study. (D) *Bt*miR29-b level was significantly higher in *Bt*miR29-b-overexpressed tomato plants. (E) The *Bt*miR29-b level decreased upon whitefly infestation in amiR29-b-expressed tomato plants. Values are mean ± SEM; $n = 6$ (3 tomato plants for each repeat). Student's *t*-test (two-tailed) was used for significant difference analysis. **, $P < 0.01$.
